# *de novo* variant calling identifies cancer mutation profiles in the 1000 Genomes Project

**DOI:** 10.1101/2021.05.27.445979

**Authors:** Jeffrey K. Ng, Pankaj Vats, Elyn Fritz-Waters, Stephanie Sarkar, Eleanor I. Sams, Evin M. Padhi, Zachary L. Payne, Shawn Leonard, Marc A. West, Chandler Prince, Lee Trani, Marshall Jansen, George Vacek, Mehrzad Samadi, Timothy T. Harkins, Craig Pohl, Tychele N. Turner

## Abstract

Detection of *de novo* variants (DNVs) is critical for studies of disease-related variation and mutation rates. We developed a GPU-based workflow to rapidly call DNVs (HAT) and demonstrated its effectiveness by applying it to 4,216 Simons Simplex Collection (SSC) whole-genome sequenced parent-child trios from DNA derived from blood. In our SSC DNV data, we identified 78 ± 15 DNVs per individual, 18% ± 5% at CpG sites, 75% ± 9% phased to the paternal chromosome of origin, and an average allele balance of 0.49. These calculations are all in line with DNV expectations. We sought to build a control DNV dataset by running HAT on 602 whole-genome sequenced parent-child trios from DNA derived from lymphoblastoid cell lines (LCLs) from the publicly available 1000 Genomes Project (1000G). In our 1000G DNV data, we identified 740 ± 967 DNVs per individual, 14% ± 4% at CpG sites, 61% ± 11% phased to the paternal chromosome of origin, and an average allele balance of 0.41. Of the 602 trios, 80% had > 100 DNVs and we hypothesized the excess DNVs were cell line artifacts. Several lines of evidence in our data suggest that this is true and that 1000G does not appear to be a static reference. By mutation profile analysis, we tested whether these cell line artifacts were random and found that 40% of individuals in 1000G did not have random DNV profiles; rather they had DNV profiles matching B-cell lymphoma. Furthermore, we saw significant excess of protein-coding DNVs in 1000G in the gene *IGLL5* that has already been implicated in this cancer. As a result of cell line artifacts, 1000G has variants present in DNA repair genes and at Clinvar pathogenic or likely-pathogenic sites. Our study elucidates important implications of the use of sequencing data from LCLs for both reference building projects as well as disease-related projects whereby these data are used in variant filtering steps.

## INTRODUCTION

*de novo* variants (DNVs) are important for assessing mutation rates ^1^ and have been shown to contribute to human disease (e.g., autism ^2–10^, epilepsy ^11,12^, intellectual disability ^13–16^, congenital heart disorders ^17–19^). Typically, the calling of DNVs from raw sequence data to final calls can take days to weeks. Multiple DNV workflows exist that primarily rely on CPU-based approaches ^2–7,9,10,12–15,17,20–31^. These workflows employ different strategies including strict filtering, utilizing multiple variant callers as opposed to using only one, machine-learning, and incorporation of genotypic information at other sites around the genome. Overall, there is no community consensus on a standard method for detecting DNVs. It is imperative that this process be streamlined and flexible to enable broad adoption across the community. In this study, we developed a rapid workflow to accelerate DNV calling using graphics processing units (GPUs) that is integrated into NVIDIA Parabricks ^32^ software. We also developed an equivalent, freely available open-source, CPU-based version of the workflow. Together, the GPU-based workflow, Hare, and the CPU-based workflow, Tortoise, make up HAT.

Our desire for a standardized, rapid DNV workflow stems from our interest in detecting these DNVs in the large number of whole-genome sequencing (WGS) data in families with neurodevelopmental disorders that has recently become available (https://anvilproject.org/data). Studies assessing individuals with WGS data based on DNA derived from blood have provided the field with our best estimates of DNV characteristics in humans ^1^. One recent dataset, with DNA derived from blood, consisting of 4,216 parent-child whole-genome sequenced trios from the Simons Simplex Collection (SSC) has been extensively studied for DNVs ^6,33–35^. We processed this data with HAT and found that our method performed well.

This led us to assess the newly generated, publicly available, WGS dataset from a cohort called the 1000 Genomes Project (1000G), where our initial goal was to build a control DNV dataset. Overall, 1000G is a data resource for the study of genetic variation that includes individuals from diverse genetic ancestries ^36,37^ Represented in the data are 602 trios from 18 worldwide populations (Figure S1). Moreover, as a field standard, 1000G has been utilized in many applications as a control resource for filtering of genetic variation by allele frequency and/or variant presence-absence in the dataset^38^.

One complicating factor of DNV assessment in this resource is the fact that sequencing data is generated from DNA isolated from lymphoblastoid cell lines (LCLs) ^37^ as opposed to primary tissue. Epstein-Barr Virus is used to make these LCLs and passaging over time enables the accumulation of cell line artifacts. These artifacts can complicate variant filtration schemes and the utility of this data as a frequency control. As opposed to a random accumulation of mutations in each individual, we found that 80% of 1000G individuals had an excess of DNVs and 40% of all 1000G individuals had a profile matching a B-cell lymphoma. The similarity to this cancer is problematic, and it would be imperative that this data not be used as a control in the context of the study of these and related cancers. A secondary consequence of the excess DNVs is their presence at disease-related sites whereby simple filtering schemes may accidentally remove sites of interest in patients due to their presence in 1000G.

## RESULTS

### Rapid DNV calling with GPUs

HAT consists of three main steps: GVCF generation, family-level genotyping, and filtering of variants to get final DNVs. We utilized existing features of the NVIDIA Parabricks software for rapid GVCF generation from GPU accelerated versions of GATK ^39^ HaplotypeCaller and Google DeepVariant ^40^. The run times for GVCF generation are ~40 minutes per sample on a 4 GPU node and can be run in parallel on all three family members in the parent-child trio. Post-GVCF generation, the trio is genotyped using the GLnexus joint genotyper ^41^. Finally, our post-genotyping custom DNV filtering workflow runs in ~1 hour with speedups at all steps with parallelization providing a clear advantage over CPU-based approaches (Figure 1A).

**Figure 1:**
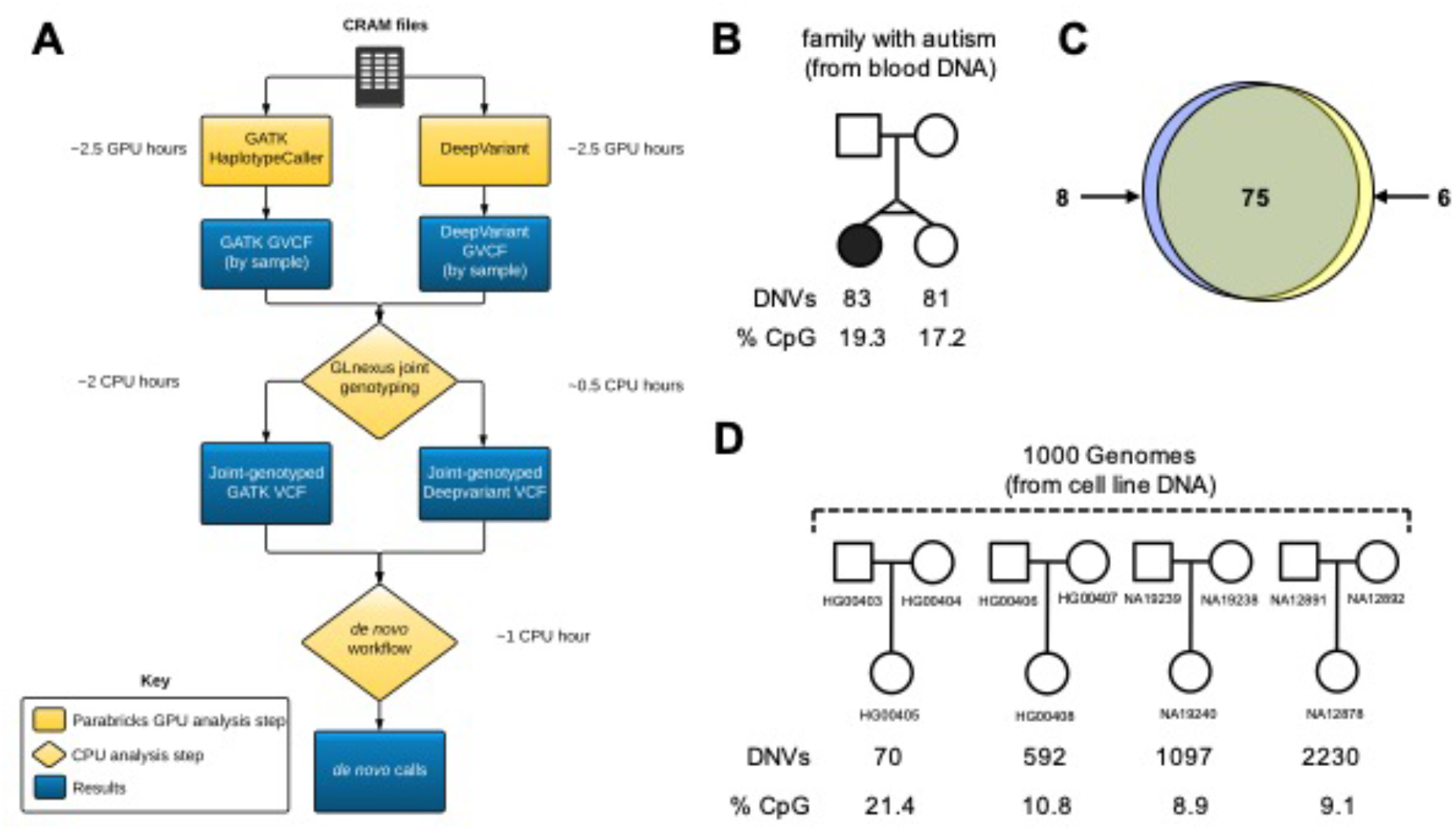
*de novo* variant calling in short-read whole-genome sequencing data. A) *de novo* workflow for detection of *de novo* variants (DNVs) from aligned read files (crams); B and C) Benchmarking DNV workflow in a monozygotic twin pair sequenced from DNA derived from blood; D) DNV detecting in four trios in the 1000 Genomes Project.

To benchmark HAT, we tested it on a monozygotic twin pair with WGS data derived from blood DNA. These individuals should share the same DNVs from generation in the germline. However, they may differ at some sites if DNVs occur in a post-zygotic, somatic manner. The twins shared 75 autosomal DNVs and contained 83 and 81 autosomal DNVs, respectively (Figures 1B and 1C). The percent CpG was 19.3% and 17.2%, respectively and in line with previous published estimates of ~20%^1,6^ (Figures 1B and 1C). As this monozygotic twin pair was discordant for the phenotype of autism, we also tested whether there were any protein-coding DNV differences between the two twins. These would potentially be relevant for autism, but there were no such differences.

To establish a DNV callset from the 1000G data as a control, we started with the assessment of DNVs with HAT in four trios from the 1000G (Figure 1D). Two were chosen at random (i.e., HG00405, HG00408) and two were chosen because they were “famous” trios assessed in many other studies (i.e., NA12878 ^23,42^, NA19240 ^23^). One of these trios (HG00405) had 70 DNVs and a CpG percent of 21.4 as we would have expected from DNA derived from blood. To our surprise, the other trios had varying numbers of DNVs from 592 to 2,230 with NA12878 (arguably the most studied individual in 1000G) having the most DNVs. With the increase in DNVs the CpG percent dropped considerably down to ~10%. We also assessed 3,598 of the DNVs from the four trios by manual visual inspection of the underlying reads in each family member (Table S1) and found that 93.6% of the variants appeared to be true DNVs, 4.9% were inherited, and 1.5% were low confidence calls.

### Differences in DNVs in blood and LCLs

Our initial observations led us to focus on two main cohorts: the 602 trios from 1000G (Table S2) with DNA derived from LCLs and 4,216 trios from the Simons Simplex Collection (SSC) with DNA derived from blood. In the 1000G data we detected 445,711 total DNVs in the cohort (Table S3). There were 740 ± 968 DNVs per individual (Table S4) with a clear bimodal distribution (Hartigan’s dip test: D = 0.033, p-value = 1.32 × 10^-4^) wherein some individuals contained an excess of DNVs (Figure 2A). In the SSC, we identified 329,589 total DNVs in the cohort. There were 78 ± 15 DNVs per individual (Figure 2B, Table S5). The values derived from the SSC data are in line with expectation and highlight the effectiveness of our DNV workflow. However, the values in the 1000G are higher than expected and we estimated the number of individuals with appropriate numbers of DNVs by splitting the 1000G data into two groups: individuals having less than or equal to 100 DNVs (n = 123) and individuals with greater than 100 DNVs (n = 479). This estimate suggests that only 20.4% of trios in the 1000G have the correct number of DNVs and we thought those with excess DNVs may have cell line artifacts due to culturing of LCLs.

**Figure 2:**
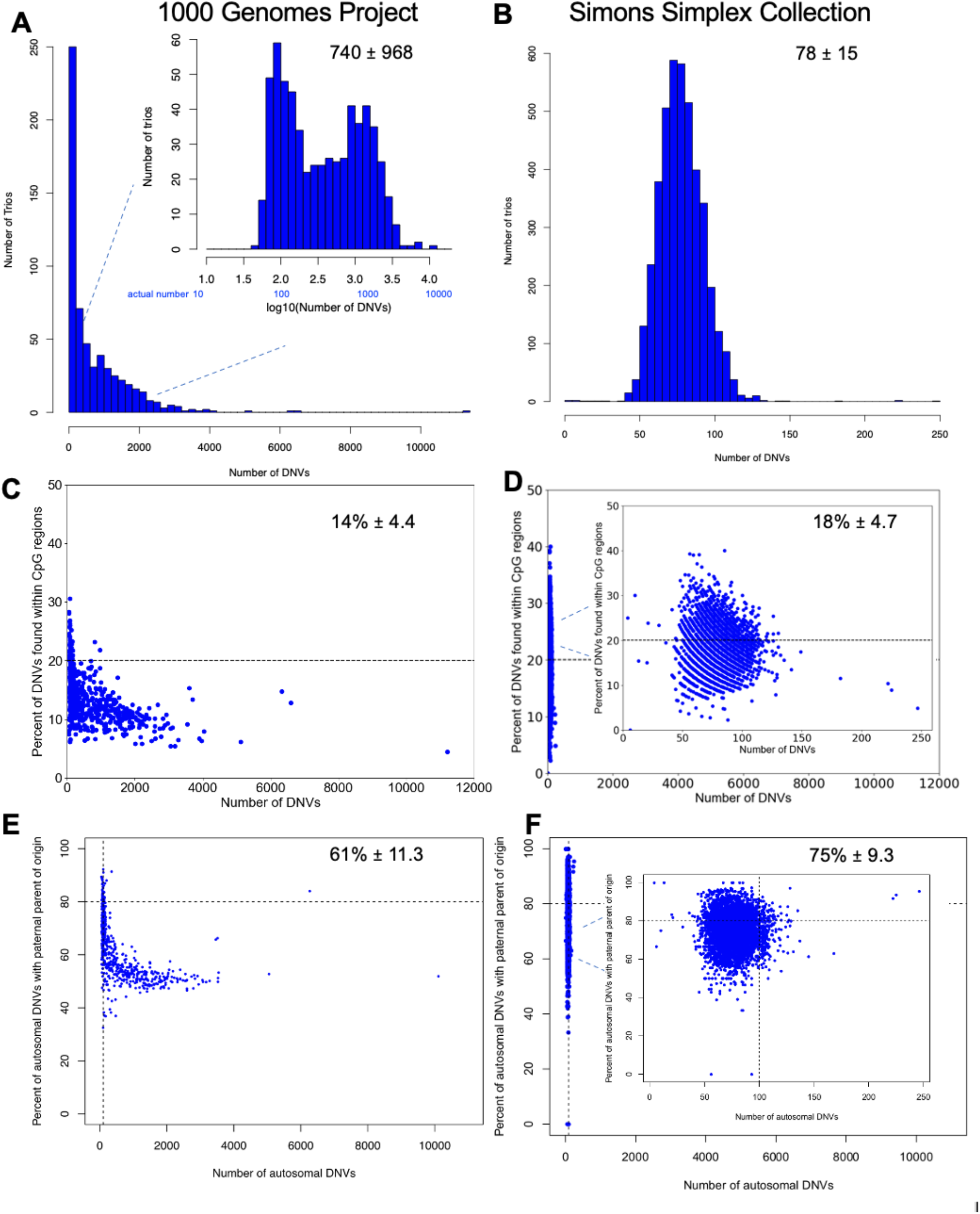
Comparison of characteristics of DNVs detected in1000 Genomes Project (1000G) and Simons Simplex Collection (SSC) callsets. A) Histogram of DNV counts from 1000G in 602 trios; B) Histogram of DNV counts from SSC in 4216 trios; C); C) Percent of DNVs found within CpG sites versus the total number of DNVs for 1000G; D) Percent of DNVs found within CpG sites versus the total number of DNVs found for SSC; E) Percent of autosomal DNVs with paternal parent of origin versus the total number of DNVs for 1000G; F) Percent of autosomal DNVs with paternal parent of origin versus the total number of DNVs for SSC.

We assessed two main features of typical DNVs to investigate our hypothesis that the excess DNVs found in individuals were cell line artifacts. These features are DNVs at CpG locations and the percent of DNVs arising on the paternal chromosome. As mentioned previously, the percent of DNVs at CpG should be ~20% and the percent of DNVs arising on the paternal chromosome should be ~80%^43^. We saw that in the 1000G trios 14 ± 4.4% of DNVs per individual occurred at CpG sites (Figure 2C) with individuals with less than or equal to 100 DNVs having17.4 ± 5.2% DNV at CpG and in families with greater than 100 DNVs 12.7 ± 3.6% DNV at CpG. The difference in DNVs at CpG sites between these two groups was significant (Wilcoxon rank sum: p-value < 2.2 × 10^-16^). In the SSC, the percent of DNVs at CpG was 18 ± 4.7%and in line with expectation (Figure 2D). In the 1000G, the percent of DNVs that were phase-able for parent-of-origin was 37.2 ± 7.5% (Figure S2). Of the phased variants, 61 ± 11.3% were on the chromosome of paternal origin (Figure 2E, Table S6). In the families with less than or equal to 100 DNVs this rose to 72.0 ± 8.5% and in the families with greater than 100 DNVs it fell to 58.6 ± 10.3%. This difference in percent phased variants of paternal origin was found to be significantly different (Wilcoxon rank sum: p-value < 2.2 × 10^-16^). The drop leveled off to ~50% in the individuals with the most DNVs (Figure 2D). In the SSC, we were able to phase 37% of DNVs (Figure S3) with the percent of DNVs phased to paternal chromosome of origin was 75% ± 9.24 and that was also in line with expectation (Figure 2F).

We also tested whether there was a difference in the allele balance (AB) in the child at DNV sites in the 1000G and the SSC (Figure S4). We found that the 1000G had a mean AB of 0.42 and in the SSC it was nearly perfect at an AB of 0.49 in line with expectation of 0.5. This indicated a lower average AB level in 1000G from newly arising mutations from cell line artifacts.

Overall, these comparisons showed that the individuals in the 1000G with less than or equal to 100 DNVs behaved more like true DNVs in regard to CpG percentage, percent arising on the paternal chromosome, and allele balance. This also was true for the SSC trios where DNA was derived from blood. However, individuals in the 1000G with > 100 DNVs did not have statistical properties of true DNVs providing evidence they may be cell line artifacts.

### DNVs by 1000G population

While we expected there to be no difference in DNV counts per individual by ancestry we sought to see if there were any populations with excess DNVs (Figure 3A). The population with the most DNVs was the CEU having on average 1,688 DNVs per individual. We hypothesized that this may be because the CEU is one of the oldest cohorts in the 1000 Genome Project dating back to the HapMap project ^44^ and these individuals may have cell lines that have been cultured more over time than other populations.

**Figure 3:**
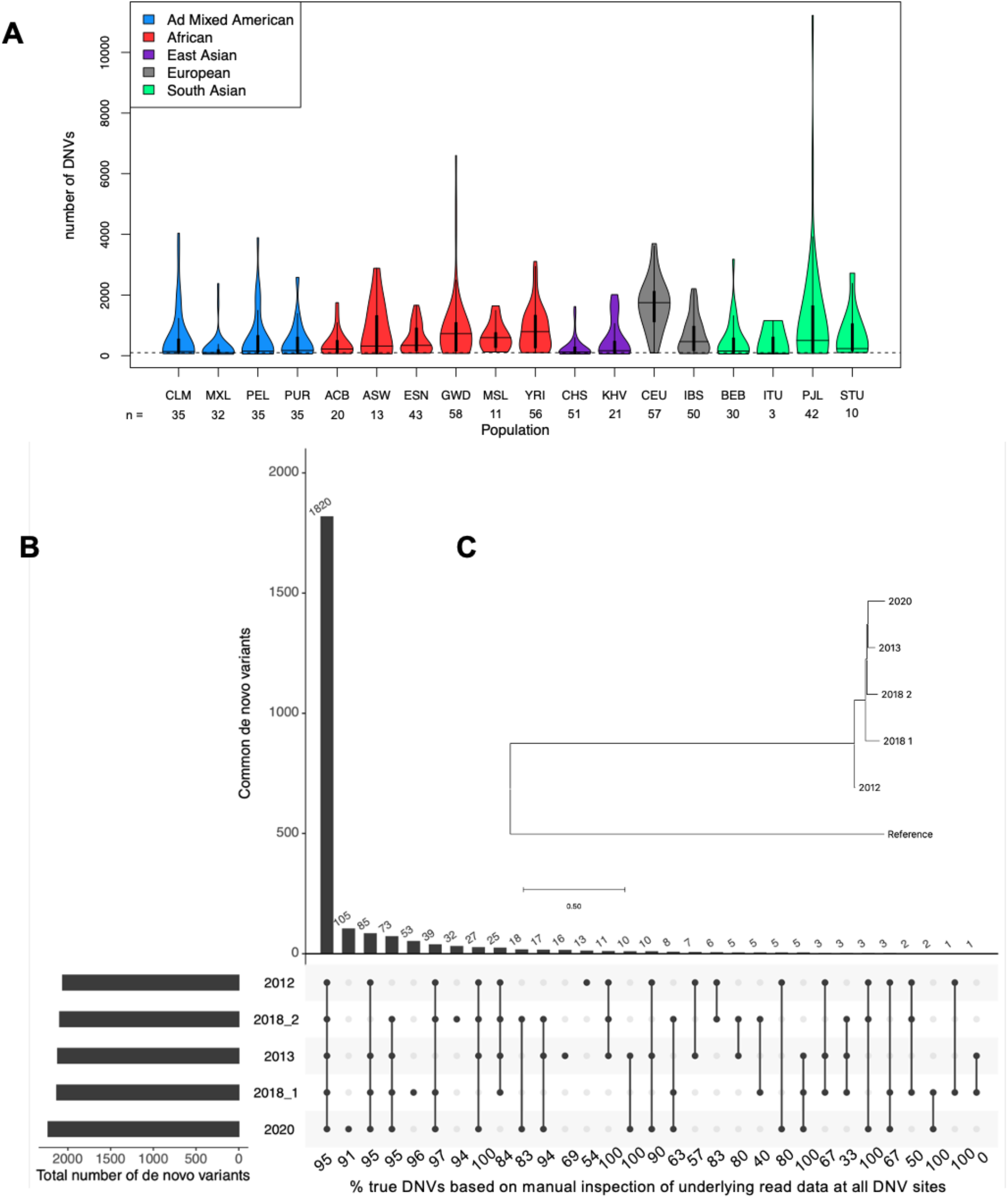
Assessment of five replicates of NA12878. A) Population distribution of 1000G dataset. B) UpSet plot demonstrating the number of variants detected in the replicates (at the bottom of the plot the percent of true DNVs is listed for each category); C) Phylogenetic tree of the five replicates.

### DNVs increase over time

We utilized the fact that the 1000G individual NA12878 has been studied and sequenced multiple times over the past ten years by WGS ^37^ (SRA identifiers: SRR944138 and SRR952827). Presumably, across time, the utilization of NA12878 has required additional culturing of this cell line, and potentially even by different laboratories. We aggregated five Illumina WGS datasets from this individual, downsampled them to ~30x coverage, and assessed them with HAT. The data for this individual ranged from the year 2012 to the year 2020 and we found that the 2012 experiment had the least DNVs (n = 2,060) and the 2020 experiment had the most DNVs (n = 2,230) (Figure 3B). Overall, the five replicates had a large overlap of DNVs (n = 1,820) across all samples. These shared DNVs constitute what were present in the ancestor of all the cell line replicates. DNVs not shared by all five replicates are sometimes shared by a subset of the replicates and are sometimes unique to the replicate. To formally assess the ancestral state, we built a phylogenetic tree based only on the DNVs and saw that the farthest replicates from each other in the tree were the 2012 and 2020 replicates (Figure 3C). To further assess the DNVs in NA12878, we randomly sampled 25 DNVs from the union dataset from the five replicates. We performed Sanger sequencing on DNA from NA12878 and her parents (NA12891, NA12892) (Figure S5-S29, Table S7). We found that 24 of the 25 DNV sites gave clear results in the Sanger sequencing with 23 confirming as real DNVs. Surprisingly, we found two sites, chr12_91353615_T_C and chr13_81142986_T_A, that were determined to be true *de novo* variants but were previously shown to be a false positive reading using Sanger sequencing ^23^. In the Sanger experiments, one site chr11_134531608_C_G showed subtle evidence for the variant allele in NA12878, so we pursued deep amplicon sequencing of this region in the trio using Oxford Nanopore Technologies (ONT) sequencing (Figure S30). This resulted in a variant allele frequency of 11% in NA12878 suggesting this is a cell line artifact. This was elevated in comparison to the background rate of 1% in NA12891 and 0% in NA12892, that is in line with expectation for error rates from ONT. Intriguingly, this DNV was only found in one of the NA12878 replicates (2018_1). Overall, this indicates that 96% of DNVs called with HAT are real (24/25) and this estimate is close to the 93.6% we saw by manual inspection of underlying read data at 3,598 DNV sites (see above).

### Genomes with cancer mutation profiles

We used mutation profile analysis ^45^ (Table S8) to determine whether the DNVs identified in individuals from the 1000G had any certain characteristics. For this analysis, we utilized a method that would enable comparisons to known mutational profiles that are either age-related (reminiscent of true DNVs) or are seen in cancers (Figure 4A and Figure 4B). There were 186 individuals (30%) that had a strong contribution of an age-related signature (Signature 1A, Signature 1B). To our surprise, the other contributing signatures in individuals were primarily those associated with B-cell lymphomas (Signature 5, Signature 9 and Signature 17) in 241 individuals (40%). This was intriguing because lymphoblastoid cell lines are generated from B-cells that are infected with Epstein Barr Virus and demonstrates that new mutations are not arising in a random manner. Rather they are being generated in a manner consistent with the development of cancer in the same cell type.

**Figure 4:**
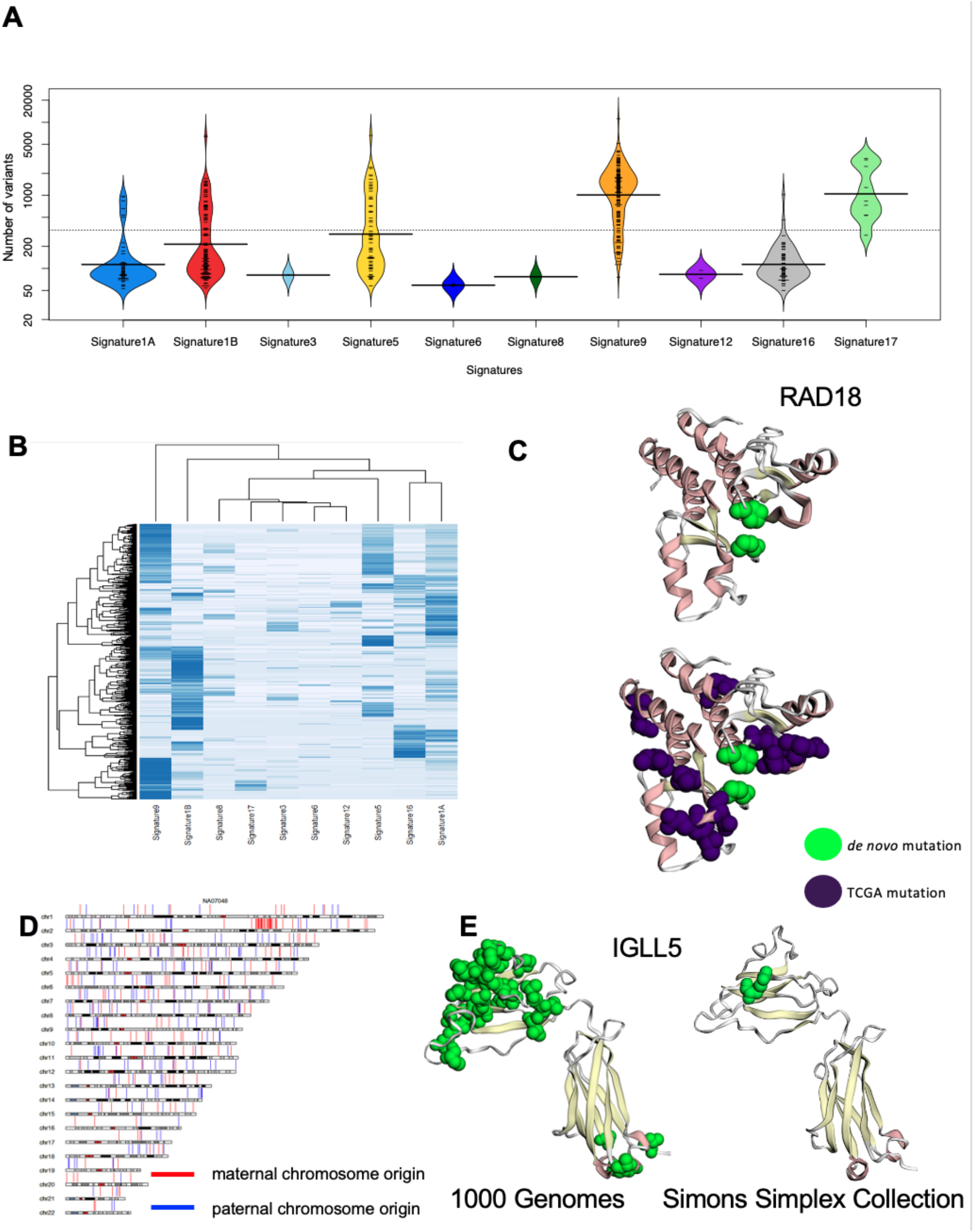
Mutational properties of DNVs. A) Mutation signature analysis showing the total number of DNVs and the individuals with each signature type; B) Heatmap of individuals based on their mutational signatures; C) Mutations in the DNA repair gene RAD18 shown on their 3D structure (and modeled using mupit). Also, shown are known cancer mutations from The Cancer Genome Atlas (TCGA); D) Location of DNVs based on their phased parent-of-origin in NA07048. Most notable there are a cluster of mutations on the maternal chromosome on chromosome 2; E) DNVs in IGLL5 shown on their 3D structure (and modeled using mupit). The image on the left is modelling variants discovered in 1000G, the image on the right is modelling variants discovered in SSC.

We further sought to determine what the mechanism was for the generation of a B-cell lymphoma-like state. First, we determined whether there was high rate of aneuploidies in the cell lines. By digital karyotyping (Table S9) we found that 595 individuals (98.8%) had a typical chromosome complement (46,XX or 46,XY), four were missing a sex chromosome (45,X0), one was 47,XXY, one had three chromosome 12 (47,XY), and one had three chromosome 9 (47,XY). This demonstrated that while these aneuploidies are occurring in some cell lines, they are probably not the main driving factor. Next, we looked at DNVs in genes involved in DNA repair and found 17 individuals contained a missense or loss-of-function in one of these genes (Table S10). Individuals with B-cell lymphoma profiles and disruptive mutations in DNA repair genes included mutations in the following genes *FANCF* (HG01126), *MUS81* (NA10838), *POLB* (NA10838), *POLD1* (NA19677), *POLE* (HG01096), *RAD18* (NA12864) (Figure 4C), *RAD51* (HG02683), *RPA4* (HG02630), and two individuals with mutations in *FANCA* (HG02841, HG03200) and *WRN* (HG04115, NA19161), respectively (Table 1). Third, we looked at Epstein Barr Virus load in each of the genomes (Table S11) and found that there was a weak, yet significant, correlation with the number of DNVs (p = 2.32 × 10^-5^, r = 0.17) (Figure S31). By visual inspection of phased variation in all individuals we also identified individuals with clusters of mutations (e.g., NA07048, Figure 4D, Figure S32).

**Table 1.**
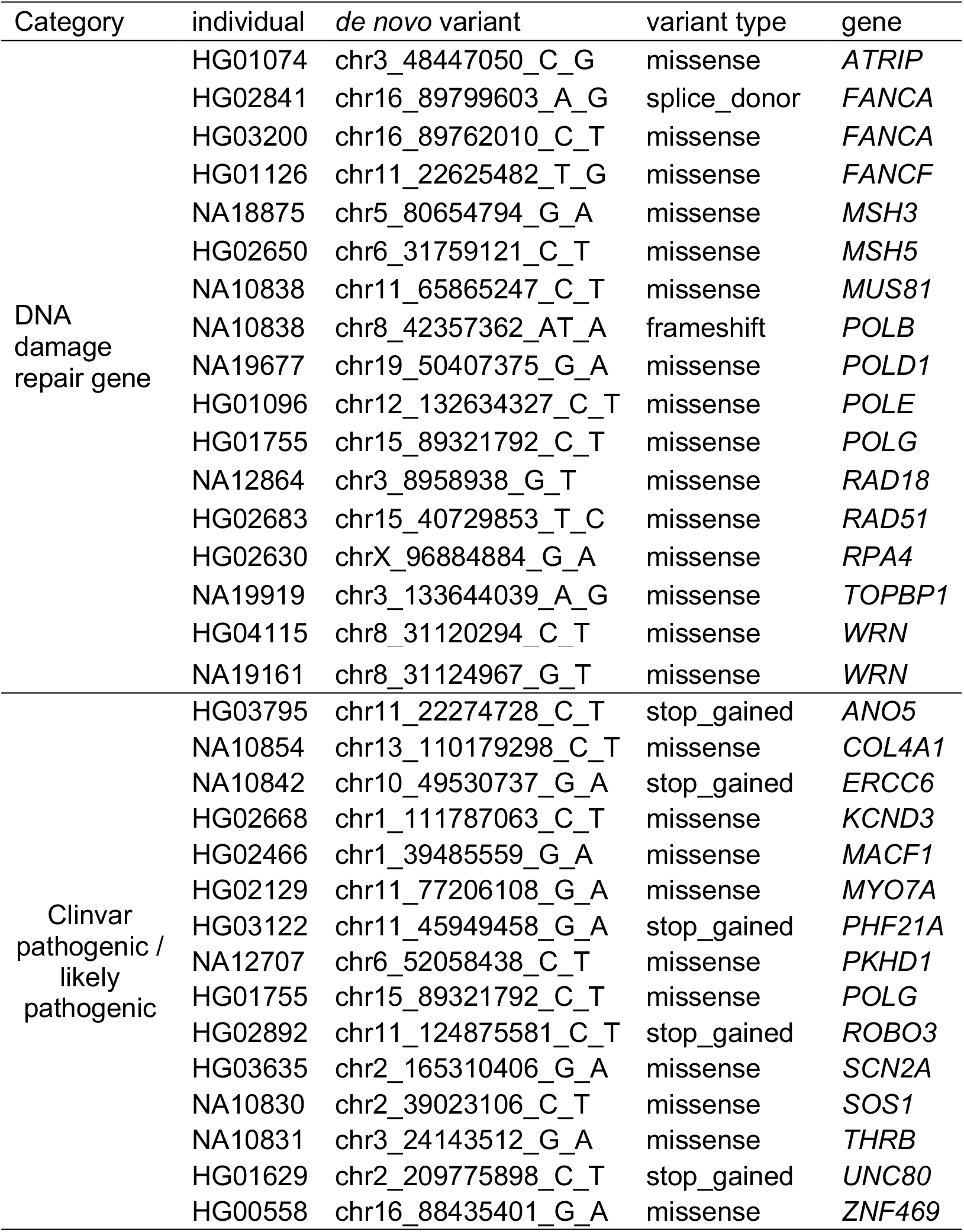
DNVs in DNA damage repair genes and clinically relevant variants

### Excess of DNVs in IGLL5

We applied a multi-phase approach to determine if there were any genes with enrichment of protein-coding DNVs in individuals with greater than 100 DNVs. In the first phase, we tested whether there was genome-wide significance for enrichment of protein-coding DNVs (missense, loss-of-function) in any specific genes. By application of two methods (chimpanzee-human, denovolyzeR), we identified 29 significant genes (*ARMC3, BCL2, BCR, C6orf15, CCDC168, CSMD3, EGR3, EXO1, HLA-B, HLA-C, IGLL5, KMT2D, LINGO2, LTB, MEOX2, MUC16, MUC22, NPAP1, PCLO, PRPF40A, RUNX1T1, SGK1, STRAP, TMEM232, TNXB, TTN, WDFY4, XIRP2, ZNF488*) with excess of DNVs (Table S11). In the second phase, we tested these 29 genes to see whether there were significantly more protein-coding DNVs in individuals with greater than 100 DNVs in comparison to individuals with less than or equal to 100 DNVs. Only *IGLL5* was significant in this comparison (1.79 × 10^-3^) (Table S12, Table S13, Figure 4E). To test whether this finding was relevant only to LCLs, we looked for protein-coding DNVs in SSC and only found one missense variant (Figure 4E). This gene did not have significant excess of DNVs in SSC.

### DNVs identified in clinically relevant variants

We tested whether any of the DNVs detected were already known to be pathogenic or likely-pathogenic in the Clinvar ^46^ database (Table 1). There were 15 mutations meeting these criteria (Table S14). We rescored these variants using Franklin software to assess their pathogenicity and found that 13 were also pathogenic or likely-pathogenic by this approach. Twelve of these variants were associated with described phenotypes in Clinvar. These included a missense variant in *SOS1* involved in Noonan syndrome, a missense variant in *SCN2A* involved in seizures, a stop gained variant in *UNC80* involved in a syndrome with hypotonia, intellectual disability, and characteristic facies, a missense variant in *THRB* involved in thyroid hormone resistance, a missense variant in *PKHD1* involved in polycystic kidney disease, a stop-gained in *ERCC6* involved in Cockayne syndrome, a stop-gained in *ANO5* involved in gnathodiaphyseal dysplasia, a stop-gained in *PHF21A* involved in inborn genetic disease, a missense in MYO7A in Usher syndrome type 1, a stop-gained in *ROBO3* in Gaze palsy with progressive scoliosis, a missense in *COL4A1* involved in inborn genetic disease, and a missense in *POLG* involved in POLG-related disorder.

## DISCUSSION

While the 1000G data has been extensively studied in the past, there has been no previous cross-cohort assessment of DNVs. This limitation is primarily because family-based sequencing was not available until 2020 when this cohort was sequenced by high-coverage short-read WGS ten years after the initial ground-breaking publication on the 1000G ^47^. Determining DNV profiles across this dataset of diverse individuals is critical for assessment of mutation rates in the human population, while also providing a more complete catalog of all genetic variants within these individuals. The decision to sequence these individuals using DNA derived from lymphoblastoid cell lines was a practical one. However, it opened the door to the possibility of cell line artifacts, while simultaneously introducing a dynamic aspect to this extensive set of controls. As control samples, the cell lines that were used as the inputs for the 1000G are still actively used across laboratories, acting as matched controls for workflows to known sets of variants. The large distribution of DNVs across the 1000G suggest that a subset of the control source inputs are dynamic, and in some cases, harbor a spectrum of genetic variants associated with B-cell lymphomas or named clinical syndromes. Laboratories using control samples from the 1000G should account for both the presence and dynamic nature of the reported DNVs and in some cases may consider changing which control samples to use within the laboratory to avoid any of the associated issues with the presence of DNVs. Additionally, other public efforts to establish reference data sets using cell lines should consider the impacts of DNVs on their project design.

We utilized a novel and accelerated analysis workflow to detect DNVs from short-read, whole-genome sequencing data. We showed this new workflow is of high-quality by running it on 4,216 trios with WGS, from the SSC, on DNA derived from blood. This analysis revealed expected number of DNVs, percent of DNVs at CpG sites, percent of DNVs phased to the paternal chromosome of origin, and average allele balance of the DNV. This was an important analysis and was in contrast to our DNV analysis of the 1000G. In total, we identified 445,711 DNVs in the 602 children from 1000G assessed in this study. We provide a cross-cohort joint-genotyped VCF, family-level VCFs, DNV calls, and phased DNV results for the 602 trios in this study as a public community resource (Globus endpoint: “Turner Lab at WashU - DNV in 1000 Genomes Paper”, direct link: https://app.globus.org/file-manager?origin_id=3eff453a-88f4-11eb-954f-752ba7b88ebe&origin_path=%2F). Originally, it was assumed that the DNVs across the 1000G would have been random and minimal, and yet only 20% of the offspring (123 children) have a number of DNVs around expectation (< 100) and the remainder have an excess of DNVs with the most extreme case being an individual (HG02683) having 11,219 DNVs. We hypothesized that the excess DNVs were cell line artifacts and found multiple lines of evidence to support this hypothesis, including a reduction in the percent of DNVs at CpG as well as the reduction in percent phased to the paternal parent-of-origin chromosome with increasing DNVs, respectively. A detailed analysis of individual NA12878, who has been studied various times over the years, revealed increasing DNVs in the more recently sequenced samples also supporting this hypothesis. The changes in the DNVs for NA12878 suggest the dynamic nature of the DNVs, demonstrating that the number is increasing over time.

When mutational signature analysis was performed on this new set of DNVs, the most common mutation signatures were those seen in B-cell lymphomas. This signature was found in 40% of individuals in the 1000G. This is important as the lymphoblastoid cell lines are generated from B-cells and points to a non-random accumulation of mutations that are in line with the development of cancer in this cell type. In particular, we identified mutations in key DNA repair genes as well as a statistically significant excess of DNVs in *IGLL5 ^48,49^*. This gene is found to be mutated in B-cell lymphomas and protein-coding DNVs are identified in 27 individuals in this cohort; all of which have >100 overall DNVs. From our work, we identify two contributing factors causing these higher levels of DNVs, one is the mutation of DNA repair genes while the second is an excess of Epstein-Barr Viral load. Future work using long-read sequencing and *de novo* assemblies will be imperative to identify complete viral integration in these genomes as integration sites can have impacts on cell line stability. One unexpected consequence of B-cell lymphoma mutation signatures in some individuals from the 1000G would be a new pathway to study the mechanisms and biology of the development of this cancer.

In addition to the DNA repair gene DNVs, we identified fifteen pathogenic or likely-pathogenic DNVs that had already been implicated in a database of clinical variation (Clinvar). This calls into question the use of the 1000G data as a control for both B-cell lymphomas and more generally for DNVs identified in clinical patients. More importantly, the extensive spectrum of DNVs that can appear in a cell line call into question the use of control samples derived from lymphoblastoid cell lines. Currently, to our knowledge the Genome in a Bottle and Human Pangenome Reference Consortium (HPRC) are building reference databases and pangenomes using DNA from lymphoblastoid cell lines. Although it does seem that the use of blood for some samples was at least initially discussed for the HPRC (https://www.genome.gov/Pages/Research/Sequencing/Meetings/HGR_Webinar_Summary_March1_2018.pdf), it does appear the project has defaulted to using lymphoblastoid cell lines. We find it is imperative that these efforts consider utilizing native DNA isolated from blood as the source or utilize a family-based design to identify and remove DNVs. In this way, the highest quality references can be built that will stand the test of time. Finally, we recommend that much like the Simons Simplex Collection, that studies assessing DNVs in individuals with a particular phenotype of interest, also sequence DNA from blood cells and not DNA post-culturing of lymphoblastoid cell lines.

## METHODS

### Software Code availability

The description for HAT (**H**are **A**nd **T**ortoise) can be found at https://github.com/TNTurnerLab/HAT.Hare, which was used for analyses in this paper are present at https://github.com/TNTurnerLab/GPU_accelerated_de_novo_workflow, v1.0. We also developed a fully open-source CPU-based version of the code that does not require the NVIDIA Parabricks license, Tortoise, and it is available at https://github.com/TNTurnerLab/Tortoise. We found that Tortoise is just as accurate as Hare, with high level of overlap between the two versions when tested on NA12878 and the monozygotic twin pair (Figure S33).

### 1000 Genomes trio whole-genome sequencing dataset

As described previously ^37^, a total of 602 trios from the 1000 Genomes Project (1000G) were whole-genome sequenced, from lymphoblastoid cell line DNA, at the New York Genome Center. We downloaded the publicly available aligned data files (crams), totaling around 27TB, onto the Washington University Information Technology’s Research Infrastructure Services (RIS), a LSF based, high compute server for further analysis described below. The download locations are described here http://ftp.1000genomes.ebi.ac.uk/vol1/ftp/data_collections/1000G_2504_high_coverage/1000G_2504_high_coverage.sequence.index and here http://ftp.1000genomes.ebi.ac.uk/vol1/ftp/data_collections/1000G_2504_high_coverage/1000G_698_related_high_coverage.sequence.index. Details of the 602 trios are found in Supplemental Table S2.

### Simons Simplex Collection whole-genome sequencing dataset

We downloaded Simons Simplex Collection whole-genome sequencing alignment files (crams) from SFARI Base using Globus, totaling around 239TB, onto the RIS. Importantly, these genomes were sequenced, from DNA derived from blood, at the New York Genome Center ^35^. We utilized the crams as the starting point for running in HAT. In total, we assessed 8,922 individuals from both quad (unaffected father, unaffected mother, one child with autism, one child without autism) and trio (unaffected father, unaffected mother, one child with autism) families resulting in a total of 4,216 parent-child sequenced trios. The following individuals were not present in the Globus link and were excluded from the study: SSC03147, SSC03138, SSC03133, SSC03146, SSC06708, SSC06703, SSC06699.

### Single-nucleotide variant and insertion/deletion calling

The NVIDIA Parabricks program version 3.0.0 was utilized to call single-nucleotide variants (SNVs) and small insertions/deletions (indels) with GATK ^39^ version 4.1.0 and Google’s DeepVariant ^40^ version 0.10 using default parameters (note for DeepVariant the model_type utilized is WGS). The reference genome utilized for these analyses was GRCh38_full_analysis_set_plus_decoy_hla.fa as the data was originally mapped to this reference genome ^37^. For each individual, a GVCF was generated for these two variant callers. The GVCFs were then genotyped, on a per trio basis, using the GLnexus ^41^ version 1.2.6 joint genotyper using prebuilt configs for each respective caller. Post-calling, we checked the counts of all variants and heterozygous variants per chromosome in each individual as a quality check (Figure S34).

### de novo variant calling

DNVs were called by identifying all putative DNVs in GATK and DeepVariant based on the parent and child genotypes, respectively. Specifically, the parent genotypes had to be homozygous for the reference allele (i.e., 0/0) and the child had to be, at a minimum, heterozygous for the alternate allele (e.g., 0/1, 1/1). DNVs identified in both GATK and DeepVariant (intersection of the two callers) were then identified and further filtering was carried out as follows: depth, in each trio member, at the DNV position had to be >= 10, the genotype quality of the DNV had to be > 20, the DNV had to have an allele balance > 0.25, and there could be no presence of the DNV allele present in any reads in the parents. Finally, we removed DNVs in low complexity regions, centromeres, and recent repeats from further analysis.

To assess the quality of our DNVs, we manually scored 3,980 sites, by visualizing the underlying read data in each trio member, with SAMtools version 1.9 tview. To score these sites, we looked at the first column (variant location in the read data as seen in tview images) of both parents and the proband sample to see what variants were present (example shown in Figure S35). If there was any variant in the first column of the mother or father, regardless of quality, that matched the main variant in the proband’s first column, then we denoted the variant as maternal, paternal, or both depending on whether it was the mother’s variant that matched the proband or the fathers or both parents. If the main variant in the first column of the parental samples did not match the proband’s variant, then we knew this sample would be a DNV, thus verifying our results.

As a second check of our DNVs, we randomly sampled 25 DNV sites in NA12878 and performed Sanger sequencing in NA12878 and parents (NA12891, NA12892). Primers were designed using Primer3Plus (https://primer3plus.com) to target each of the 25 variants. PCR reactions were run using the primers, genomic DNA for individuals NA12878 (Coriell tube label NA12878 * N44 12/02/2019), NA12891 (Coriell tube label NA12891 * H3 7/25/2019), and NA12892 (Coriell tube label NA12892 * F3 8/6/2019), and Thermo Scientific Phusion High-Fidelity PCR Master Mix with HF Buffer. All PCR products underwent PCR clean-up and Sanger sequencing through Genewiz (https://www.genewiz.com). Trace files with the Sanger sequencing data were assembled and visualized as chromatograms using Sequencher 5.4.6 (http://www.genecodes.com). For 24 of the variants, the result from Sanger sequencing was clear. However, for site chr11_134531608_C_G we saw evidence of the alternate allele at a low frequency. To test whether this signal was real, we pursued deep sequencing of the amplicon on an Oxford Nanopore Technologies (ONT) MinION sequencer as follows. PCR products for amplicon chr11_134531608_C_G, in each of the three individuals, underwent purification using the QIAquick PCR Purification Kit. A library of the purified products was prepared using the Oxford Nanopore Technologies (ONT) Rapid Barcoding Kit (SQK-RBK004). Sequencing of the library was performed using the ONT MinION sequencer and the MinKNOW software. The fastq output files containing the sequencing data for all three samples were mapped to the amplicon reference sequence using minimap2 ^50^ (version 2.21) and all had coverage depth > 100x. A bam file and indexed bam file were generated for each sample using SAMtools ^51^ (version 1.9). The bam files were then visualized using the Integrated Genomics Viewer ^52^ to determine the count of each nucleotide base at the variant position.

### Phasing of de novo variants

We utilized Unfazed version 1.0.2 (https://github.com/jbelyeu/unfazed) ^53^ to phase the *de novo* variants in our study with regard to the parent-of-origin chromosome. First, a bed file containing *de novo* variants was generated for each individual. Second, the *de novo* bed file, DeepVariant full genome trio VCF, and the alignment files for all trio members were run through Unfazed. Since Unfazed uses different approaches to phasing on the X chromosome in males and females, we only focus on phased variants on the autosomes in this study.

### NA12878 additional datasets

We identified additional high-coverage whole-genome sequencing data from NA12878 from the SRA (https://www.ncbi.nlm.nih.gov/sra) and other sources. These included SRA data SRR944138 from 2012 and SRR952827 from 2013, McDonnell Genome Institute data gerald_HFKWMDSXX and H_IJ-NA12878 both from 2018, and the high-coverage data from 2020. To avoid differences due to coverage, we downsampled all datasets to 30x using SAMtools. All data was re-mapped to build 38 using SpeedSeq ^54^ version 0.1.2 and run through the DNV workflow using the NA12891 and NA12892 parental WGS data from 2020 1000G. We again did a count check for total and heterozygous variants per chromosome (Figure S36).

### Phylogenetic tree of de novo variants

To assess the differences between different NA12878 replicates we built a multi-sequence FASTA file where each FASTA represents the aggregate of all possible DNVs identified in this individual. The specific steps to build the tree were as follows: 1) we first merged the samples together and converted the genotypes for each DNV site from 0/0 or 0/1 to the nucleotide counterparts (e.g., AA, CG, TC) for all of the NA12878 samples; 2) next we converted these genotype symbols to their IUPAC code; 3) we then collapsed the IUPAC symbols into a sequence per sample and placed them into a FASTA file. We also included a reference “sample”, which was just the reference allele at each DNV site and 4) we used MEGAX ^55^ version 10.2.4 to create a maximum likelihood phylogenetic tree.

### Mutation profile assessment

We utilized the deconstructSig ^45^ software version-1.9.0 inside of Parabricks to perform mutation signature analysis. The prominent signature was chosen for an individual and if there was not one prominent signature than the weights of two signature was equal to or greater than (>= 0.31) both signatures were represented in the tables and figures.

### Karyotype analysis

Read-depth based karyotypes were generated by assessment of the aligned sequence data. First, the number of reads per chromosome was calculated using SAMtools ^51^ in each individual. Second, the size of each chromosome was generated using the reference genome data and by removing locations of gaps from the reference. Third, the copy number of each of the chromosomes was calculated as follows: ((fold coverage per chromosome) / (fold coverage of chromosome 1))*2.

### Viral analysis

We ran SAMtools idxstats on all individuals to determine the number of mapped reads to each chromosome. We then calculated the copy number of EBV in each individual as follows: EBV copy number = ((mapped reads to EBV * 150 base pairs per read) / length of EBV) / ((mapped reads to chromosome 1 * 150 base pairs per read) / length of chromosome 1)

### DNV enrichment in genes

To test for DNV enrichment in genes we utilized two methods: chimpanzee-human and denovolyzeR. These were run as previously described ^8,56^.

### Annotation of protein-coding DNVs

We uploaded the DNV calls to the open-cravat program (https://opencravat.org/) and specifically identified Clinvar as one of the annotation categories. Rescoring of DNVs in Franklin was performed using Franklin (https://franklin.genoox.com).

## Supporting information

1000 Genomes Unfazed results

1000 Genomes DNV callset

All Supplemental Tables

High Res Main Figures and Supplemental Figures

## Author Contributions

J.K.N., T.T.H., C.P., and T.N.T. designed the study; J.K.N., P.V., E.F., S.S., E.I.S, E.M.P., Z.L.P., S.L.,, C.P., L.T., M.J., M.S., C.P. and T.N.T. performed analyses; J.N. and T.N.T. wrote the paper; and all authors reviewed and edited the paper.

## Data and materials availability

We provide family-level VCFs, DNV calls, and phased DNV results for the 602 trios in this study as a public community resource (Globus endpoint: “Turner Lab at WashU - DNV in 1000 Genomes Paper”, direct link: https://app.globus.org/file-manager?origin_id=3eff453a-88f4-11eb-954f-752ba7b88ebe&origin_path=%2F). 1000 Genomes Acknowledgement for deep coverage of the extended 3202 genomes (or subset thereof): The following cell lines/DNA samples were obtained from the NIGMS Human Genetic Cell Repository at the Coriell Institute for Medical Research: [NA06984, NA06985, NA06986, NA06989, NA06991, NA06993, NA06994, NA06995, NA06997, NA07000, NA07014, NA07019, NA07022, NA07029, NA07031, NA07034, NA07037, NA07045, NA07048, NA07051, NA07055, NA07056, NA07340, NA07345, NA07346, NA07347, NA07348, NA07349, NA07357, NA07435, NA10830, NA10831, NA10835, NA10836, NA10837, NA10838, NA10839, NA10840, NA10842, NA10843, NA10845, NA10846, NA10847, NA10850, NA10851, NA10852, NA10853, NA10854, NA10855, NA10856, NA10857, NA10859, NA10860, NA10861, NA10863, NA10864, NA10865, NA11829, NA11830, NA11831, NA11832, NA11839, NA11840, NA11843, NA11881, NA11882, NA11891, NA11892, NA11893, NA11894, NA11917, NA11918, NA11919, NA11920, NA11930, NA11931, NA11932, NA11933, NA11992, NA11993, NA11994, NA11995, NA12003, NA12004, NA12005, NA12006, NA12043, NA12044, NA12045, NA12046, NA12056, NA12057, NA12058, NA12144, NA12145, NA12146, NA12154, NA12155, NA12156, NA12234, NA12239, NA12248, NA12249, NA12264, NA12272, NA12273, NA12274, NA12275, NA12282, NA12283, NA12286, NA12287, NA12329, NA12335, NA12336, NA12340, NA12341, NA12342, NA12343, NA12344, NA12347, NA12348, NA12375, NA12376, NA12383, NA12386, NA12399, NA12400, NA12413, NA12414, NA12485, NA12489, NA12546, NA12707, NA12708, NA12716, NA12717, NA12718, NA12739, NA12740, NA12748, NA12749, NA12750, NA12751, NA12752, NA12753, NA12760, NA12761, NA12762, NA12763, NA12766, NA12767, NA12775, NA12776, NA12777, NA12778, NA12801, NA12802, NA12812, NA12813, NA12814, NA12815, NA12817, NA12818, NA12827, NA12828, NA12829, NA12830, NA12832, NA12842, NA12843, NA12864, NA12865, NA12872, NA12873, NA12874, NA12875, NA12877, NA12878, NA12889, NA12890, NA12891, NA12892]. The data were generated at the New York Genome Center with funds provided by NHGRI Grants 3UM1HG008901-03S1 and 3UM1HG008901-04S2. We are grateful to all of the families at the participating SSC sites, as well as the principal investigators (A. Beaudet, R. Bernier, J. Constantino, E. Cook, E. Fombonne, D. Geschwind, R. Goin-Kochel, E. Hanson, D. Grice, A. Klin, 25D. Ledbetter, C. Lord, C. Martin, D. Martin, R. Maxim, J. Miles, O. Ousley, K. Pelphrey, B. Peterson, J. Piggot, C. Saulnier, M. State, W. Stone, J. Sutcliffe, C. Walsh, Z. Warren, and E. Wijsman). We appreciate obtaining access to phenotypic and genetic data for the monozygotic twin pair on SFARI Base. Approved researchers can obtain the SSC population dataset described in this study (https://www.sfari.org/resource/simons-simplex-collection/) by applying at https://base.sfari.org.

