## Supplementary material for "*de novo* variant calling identifies cancer mutation profiles in the 1000 Genomes Project": High Res Main Figures and Supplemental Figures

Figure 1

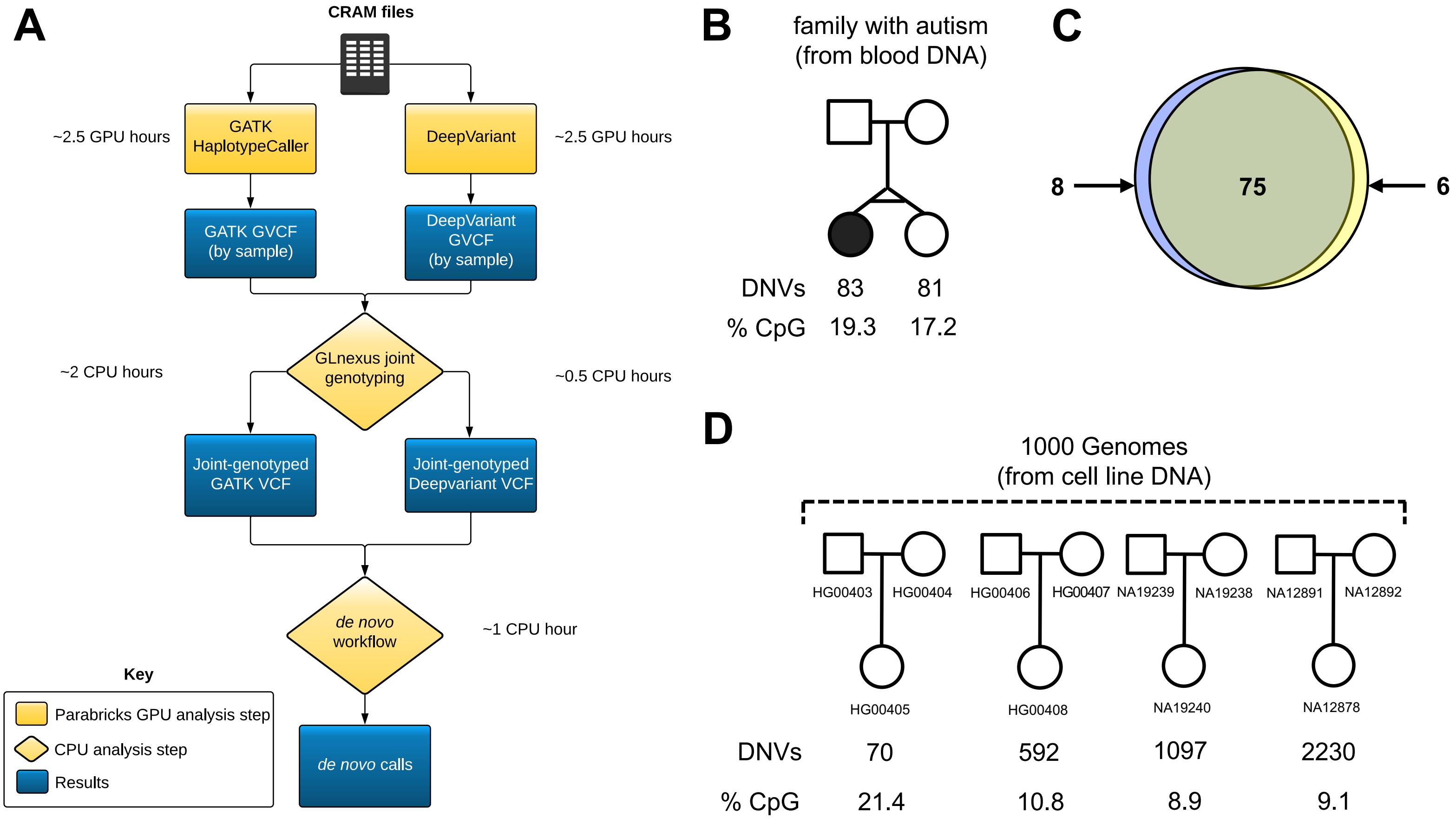
A
CRAM files
GATK HaplotypeCaller
~2.5 GPU hours
DeepVariant
~2.5 GPU hours
GATK GVCF (by sample)
DeepVariant GVCF (by sample)
GLnexus joint genotyping
~2 CPU hours
~0.5 CPU hours
Joint-genotyped GATK VCF
Joint-genotyped Deepvariant VCF
de novo workflow
~1 CPU hour
de novo calls
Key
Parabricks GPU analysis step
CPU analysis step
Results
B
family with autism (from blood DNA)
DNVs 83 81
% CpG 19.3 17.2
C
8
75
6
D
1000 Genomes (from cell line DNA)
HG00403
HG00404
HG00406
HG00407
NA19239
NA19238
NA12891
NA12892
HG00405
HG00408
NA19240
NA12878
DNVs 70 592 1097 2230
% CpG 21.4 10.8 8.9 9.1

Figure 2

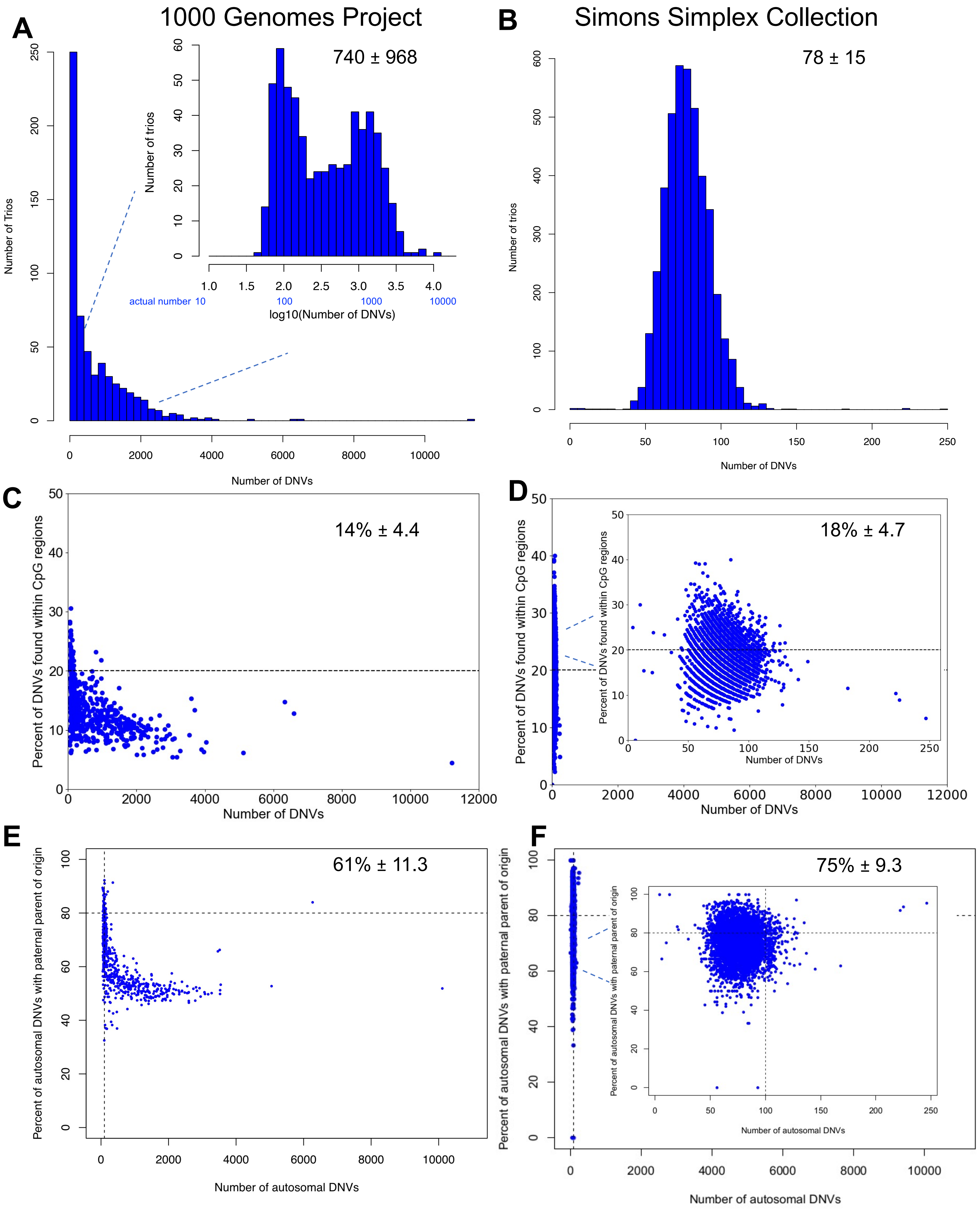

A
1000 Genomes Project
740 ± 968
Number of Trios
Number of trios
actual number 10
100
1000
10000
log10(Number of DNVs)
Number of DNVs
B
Simons Simplex Collection
78 ± 15
Number of trios
Number of DNVs
C
14% ± 4.4
Percent of DNVs found within CpG regions
Number of DNVs
D
18% ± 4.7
Percent of DNVs found within CpG regions
Number of DNVs
E
61% ± 11.3
Percent of autosomal DNVs with paternal parent of origin
Number of autosomal DNVs
F
75% ± 9.3
Percent of autosomal DNVs with paternal parent of origin
Number of autosomal DNVs

Figure 3

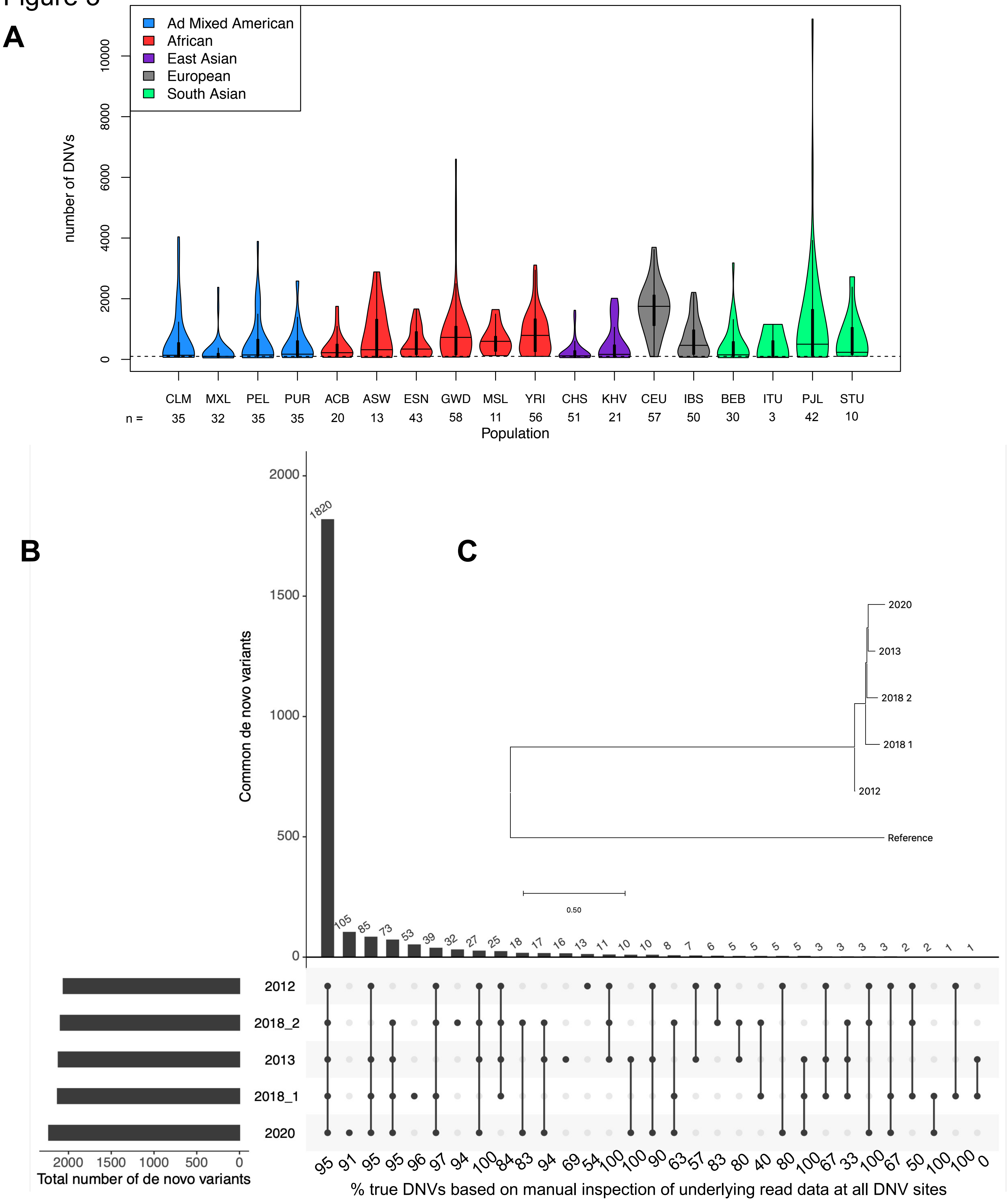
A
Ad Mixed American
African
East Asian
European
South Asian
number of DNVs
0
2000
4000
6000
8000
10000
CLM MXL PEL PUR ACB ASW ESN GWD MSL YRI CHS KHV CEU IBS BEB ITU PJL STU
n = 35 32 35 35 20 13 43 58 11 56 51 21 57 50 30 3 42 10
Population
B
C
Common de novo variants
0
500
1000
1500
2000
1820 105 85 73 53 39 32 27 25 18 17 16 13 11 10 10 8 7 6 5 5 5 5 3 3 3 3 2 2 1 1
2020
2013
2018 2
2018 1
2012
Reference
0.50
2012
2018_2
2013
2018_1
2020
2000 1500 1000 500 0
Total number of de novo variants
95 91 95 95 96 97 94 100 84 83 94 69 54 100 100 90 63 57 83 80 40 80 100 67 33 100 67 50 100 100 0
% true DNVs based on manual inspection of underlying read data at all DNV sites

Figure 4

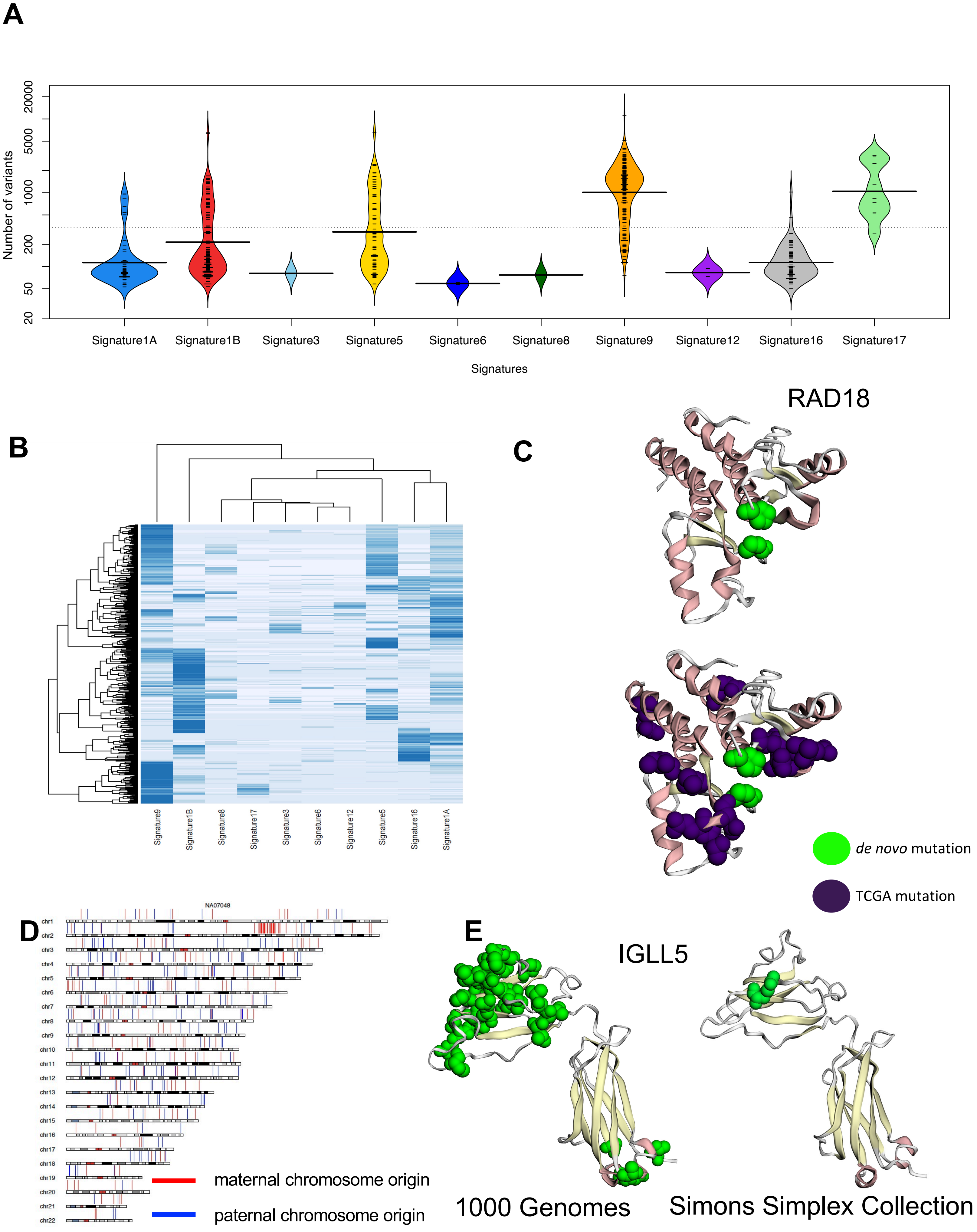
A
Number of variants
20000
5000
1000
200
50
20
Signature1A
Signature1B
Signature3
Signature5
Signature6
Signature8
Signature9
Signature12
Signature16
Signature17
Signatures
RAD18
B
Signature9
Signature1B
Signature8
Signature17
Signature3
Signature6
Signature12
Signature5
Signature16
Signature1A
C
de novo mutation
TCGA mutation
NA07048
D
chr1
chr2
chr3
chr4
chr5
chr6
chr7
chr8
chr9
chr10
chr11
chr12
chr13
chr14
chr15
chr16
chr17
chr18
chr19
chr20
chr21
chr22
maternal chromosome origin
paternal chromosome origin
E
IGLL5
1000 Genomes
Simons Simplex Collection

### Figure S1

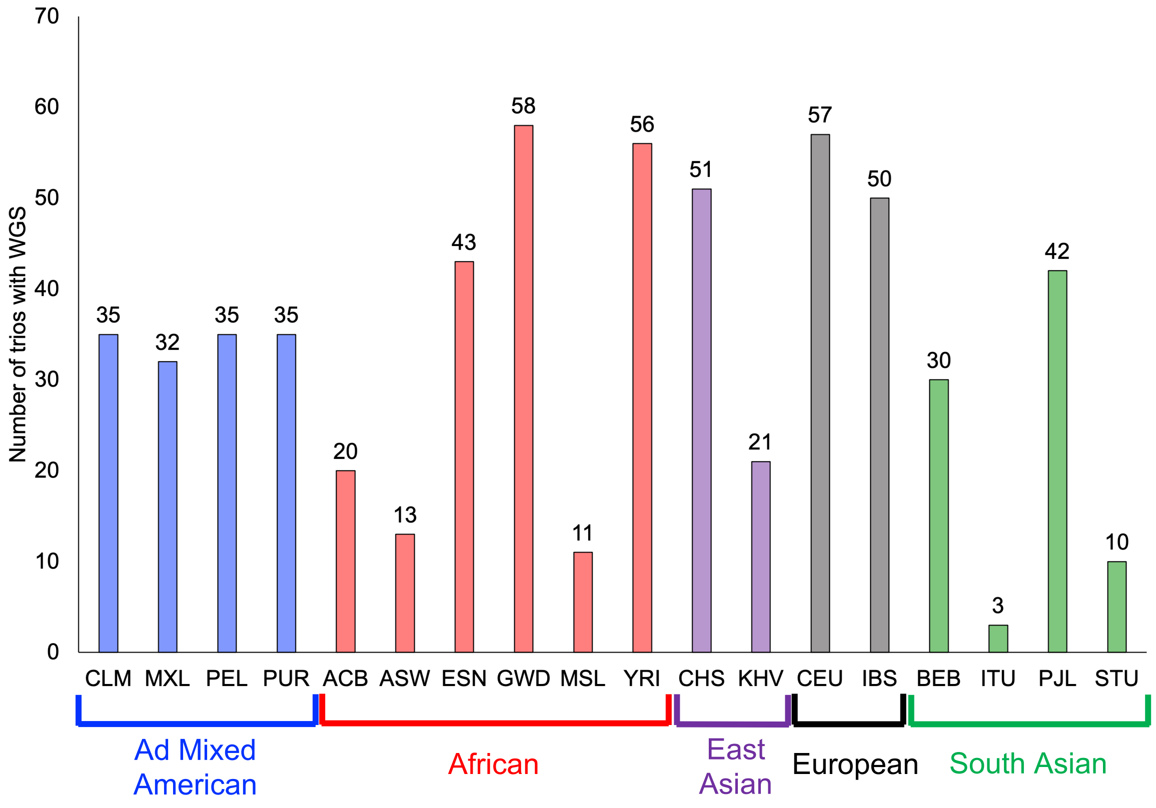

Number of trios with WGS
70
60
50
40
30
20
10
0
35
32
35
35
20
13
43
58
11
56
51
21
57
50
30
3
42
10
CLM
MXL
PEL
PUR
ACB
ASW
ESN
GWD
MSL
YRI
CHS
KHV
CEU
IBS
BEB
ITU
PJL
STU
Ad Mixed American
African
East Asian
European
South Asian

Figure S2

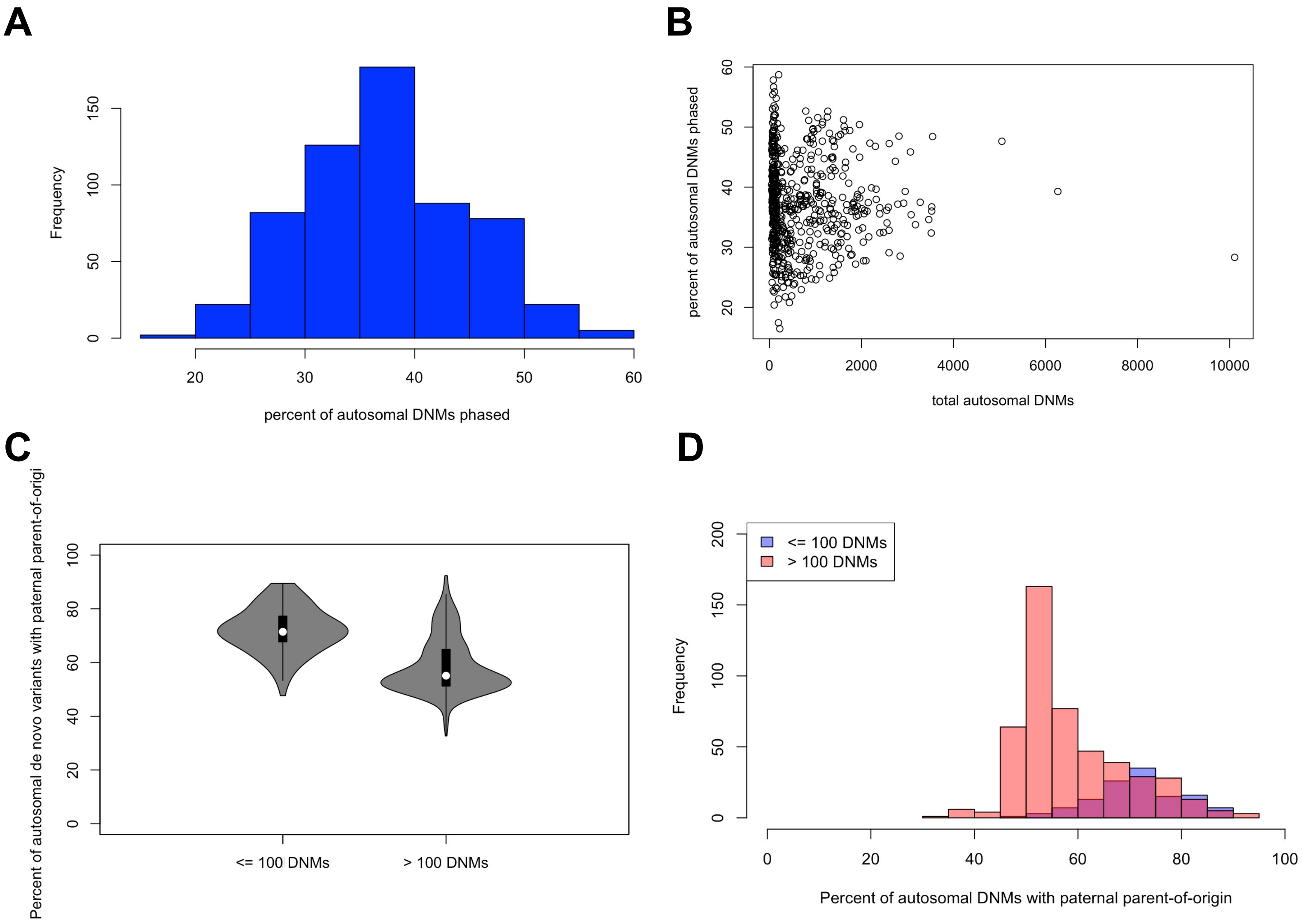
A
Frequency
percent of autosomal DNMs phased
B
percent of autosomal DNMs phased
total autosomal DNMs
C
Percent of autosomal de novo variants with paternal parent-of-origi
<= 100 DNMs
> 100 DNMs
D
<= 100 DNMs
> 100 DNMs
Frequency
Percent of autosomal DNMs with paternal parent-of-origin

Figure S3

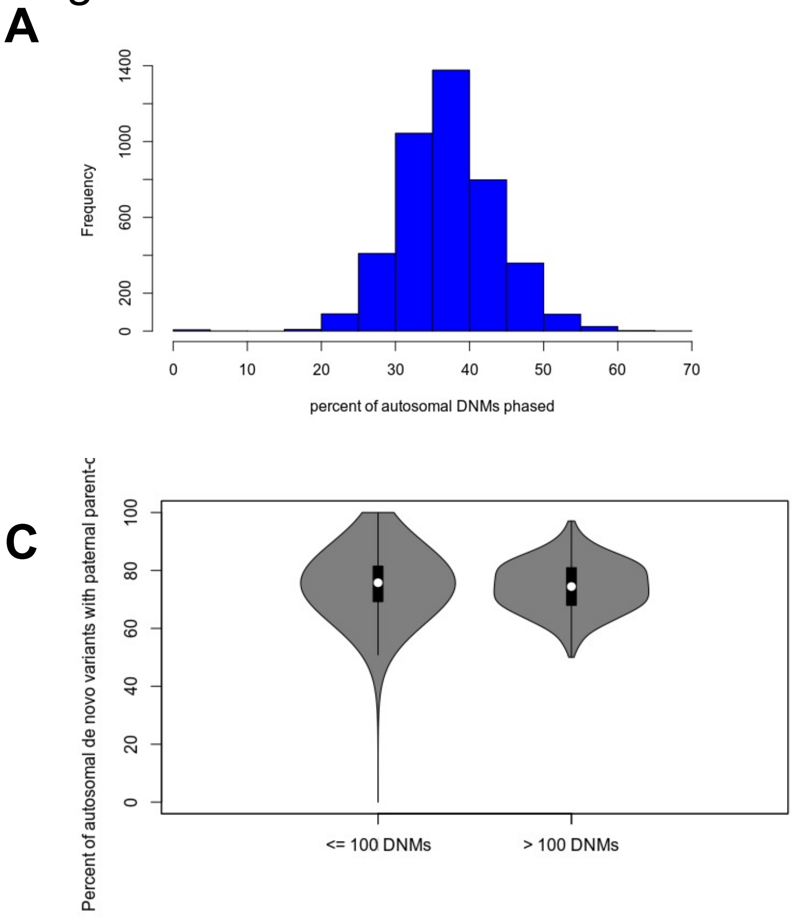

A
Frequency
percent of autosomal DNMs phased
B
Percent of autosomal DNMs phased
total autosomal DNMs
C
Percent of autosomal de novo variants with paternal parent-o
<= 100 DNMs
> 100 DNMs
D
<= 100 DNMs
> 100 DNMs
Frequency
Percent of autosomal DNMs with paternal parent-of-origin

### Figure S4

**A**

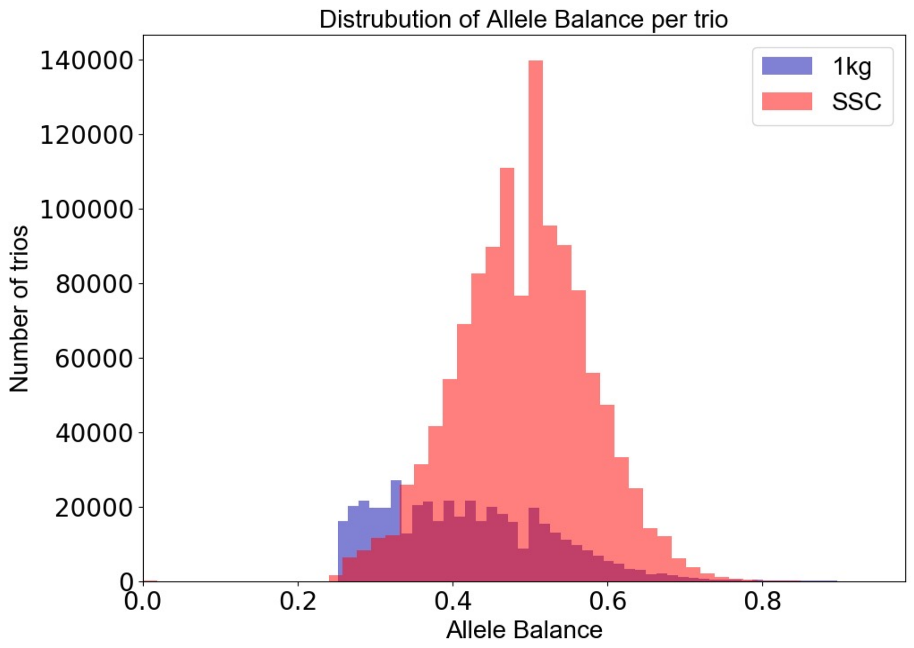


Mean for 1000 Genomes: 0.419

Mean for SSC: 0.491

### Figure S5

chr4_187891282_G_T

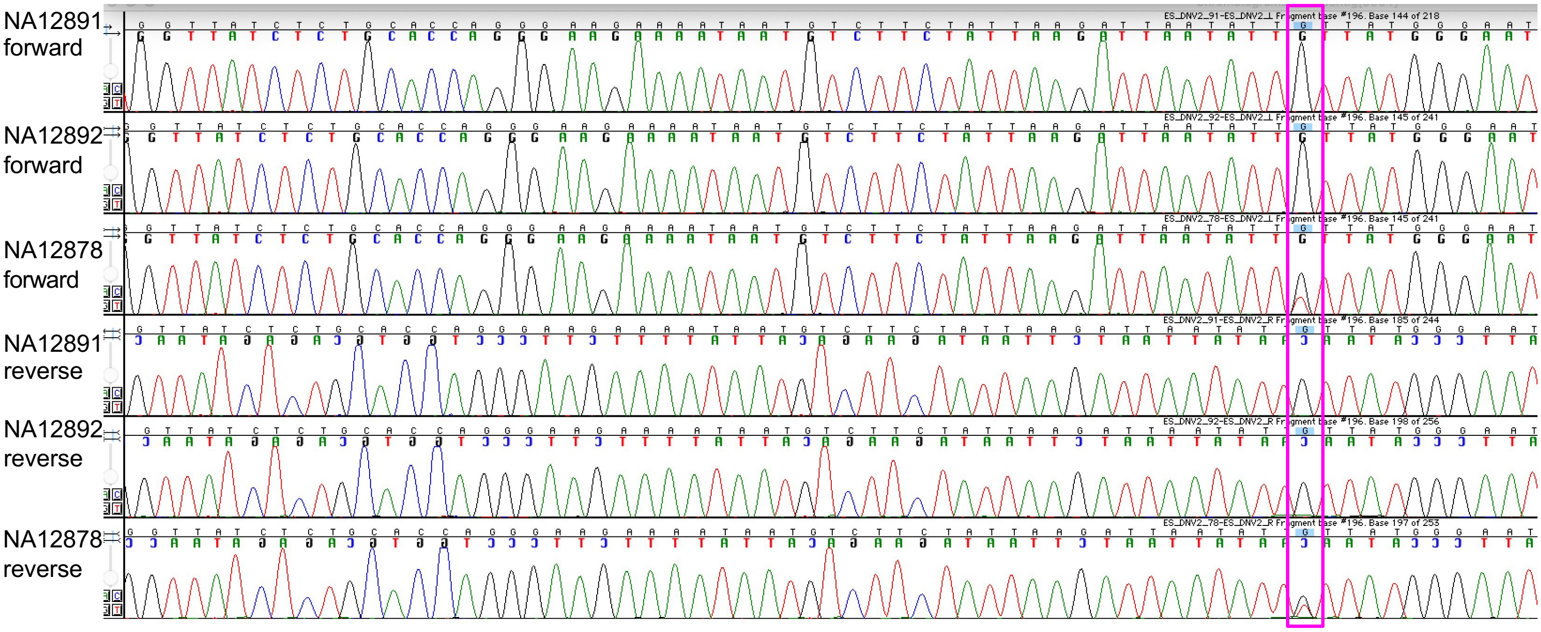

### Figure S6

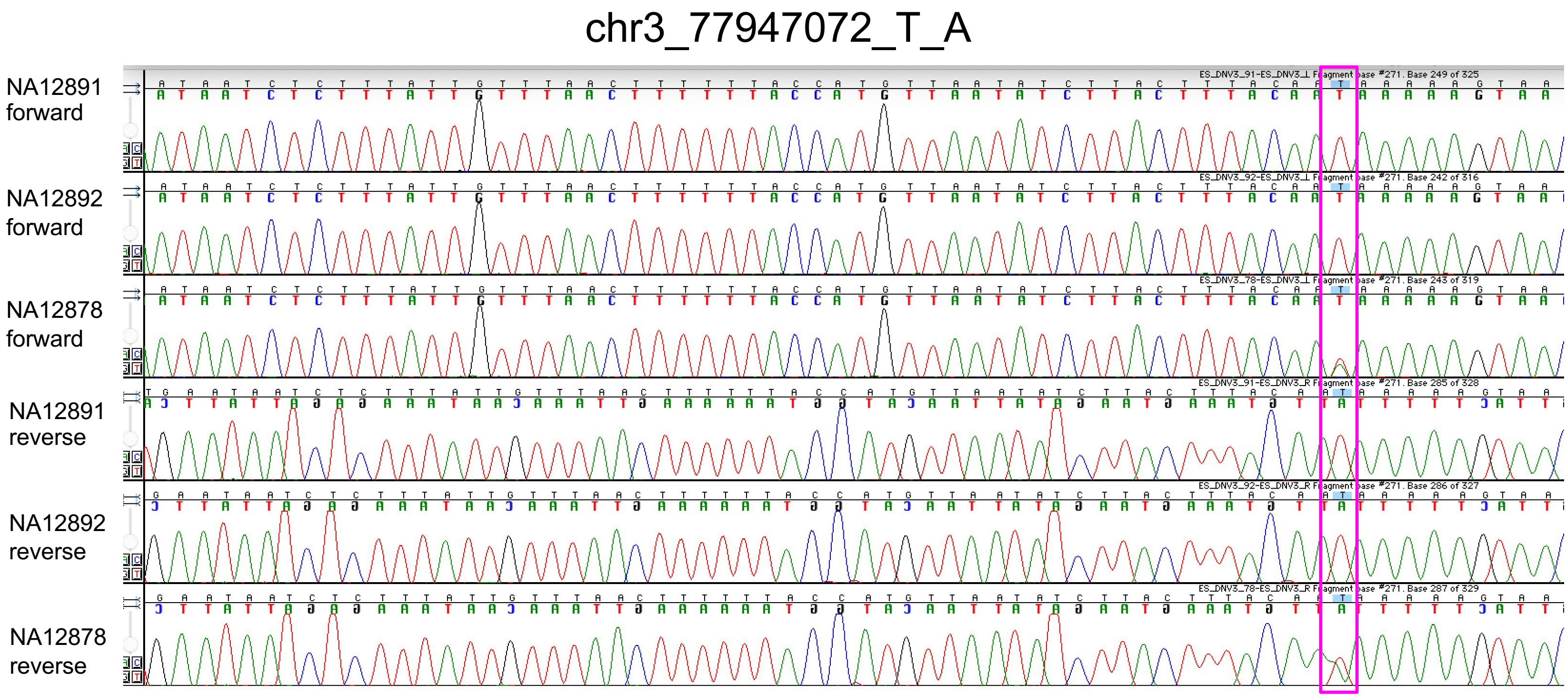
chr3_77947072_T_A
NA12891 forward
NA12892 forward
NA12878 forward
NA12891 reverse
NA12892 reverse
NA12878 reverse

Figure S7

chr8_64304207_C_A

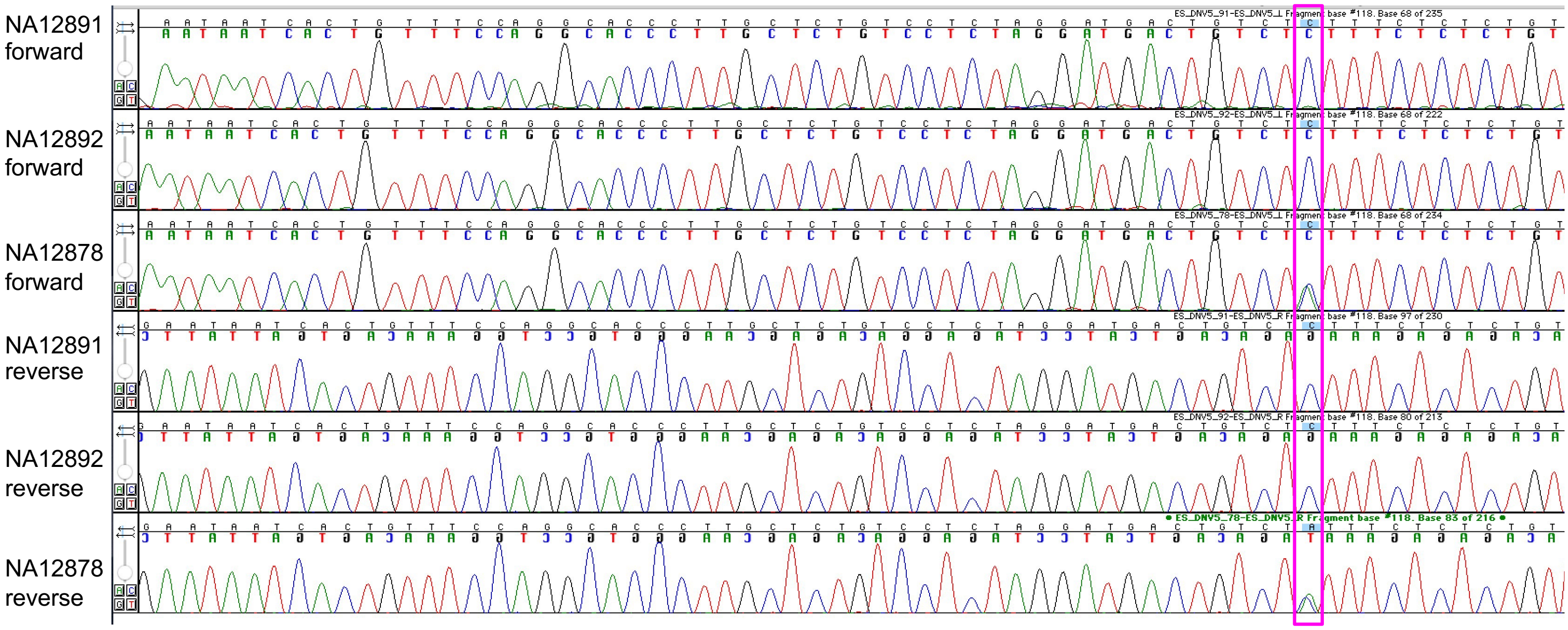

### Figure S8

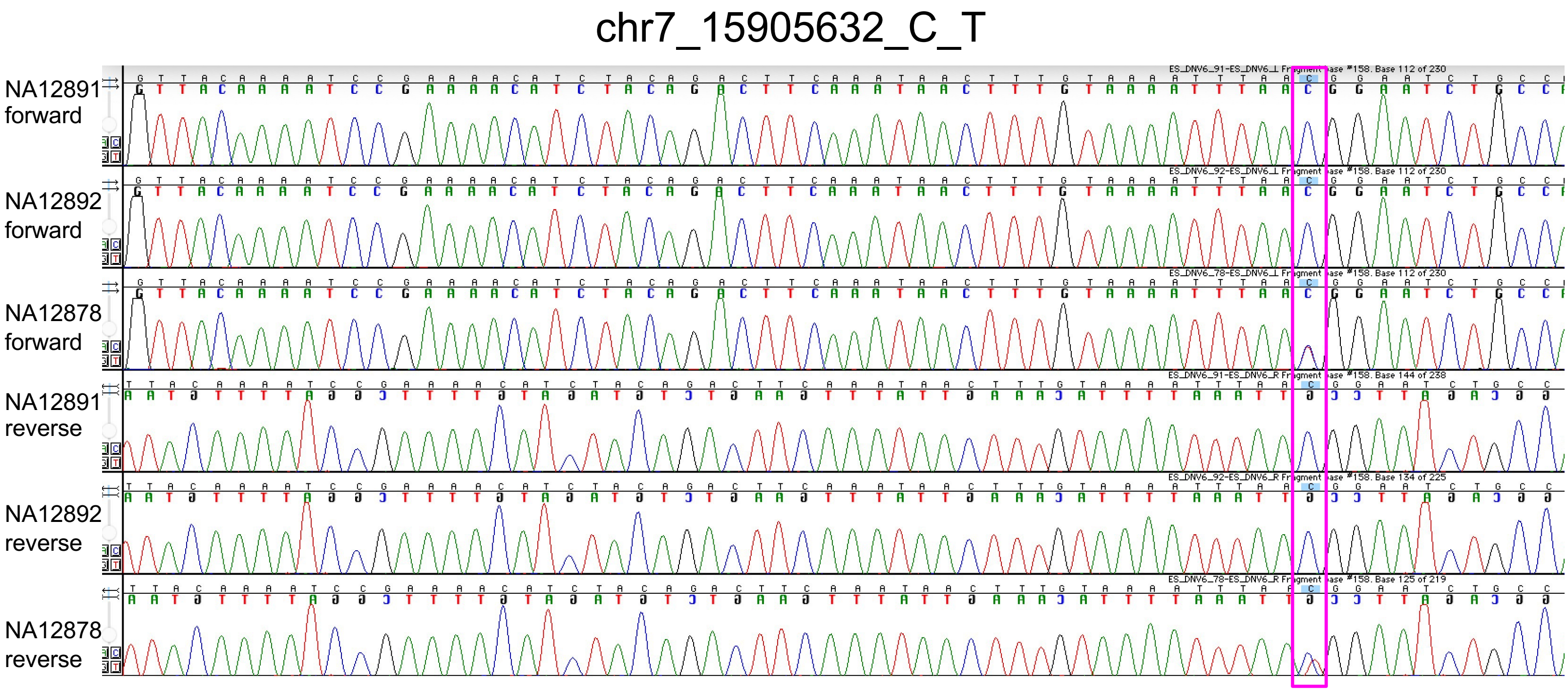

chr7_15905632_C_T
NA12891 forward
NA12892 forward
NA12878 forward
NA12891 reverse
NA12892 reverse
NA12878 reverse

### Figure S9

chr3_151755926_G_T

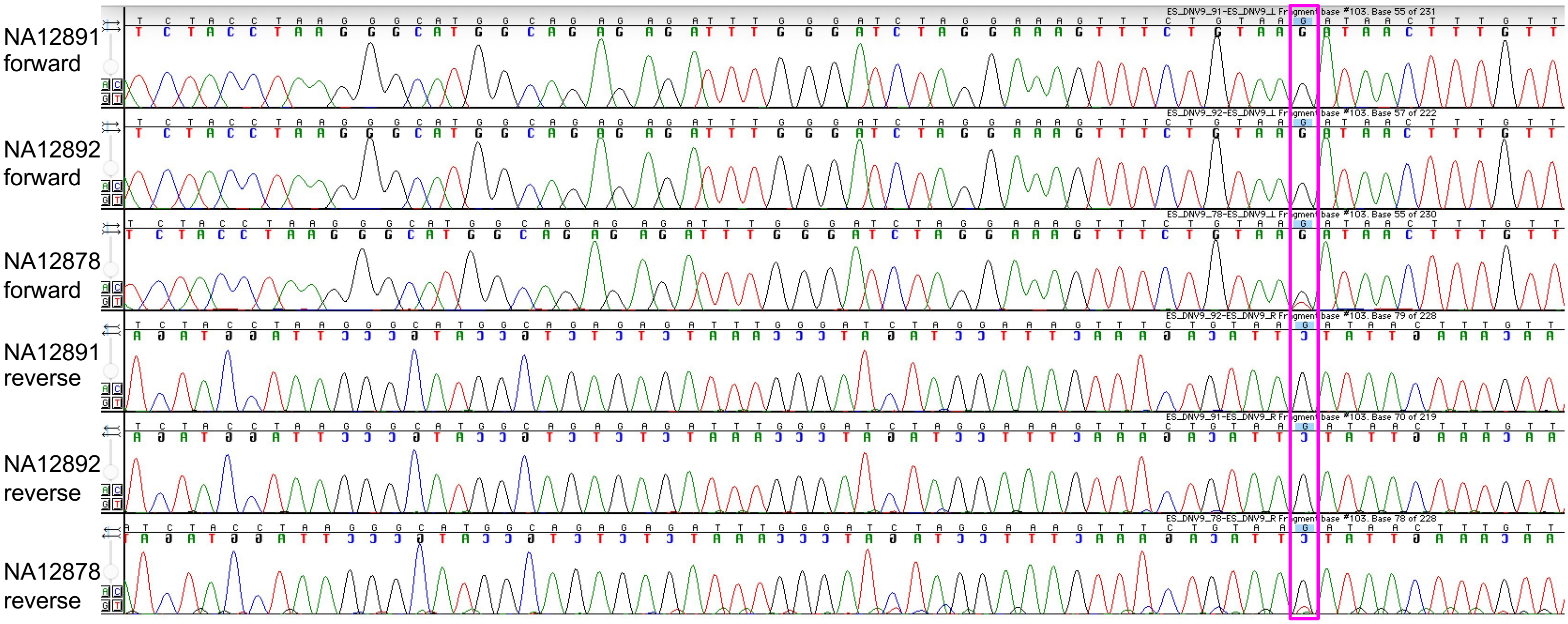

Figure S10

chr6_93124594_T_C

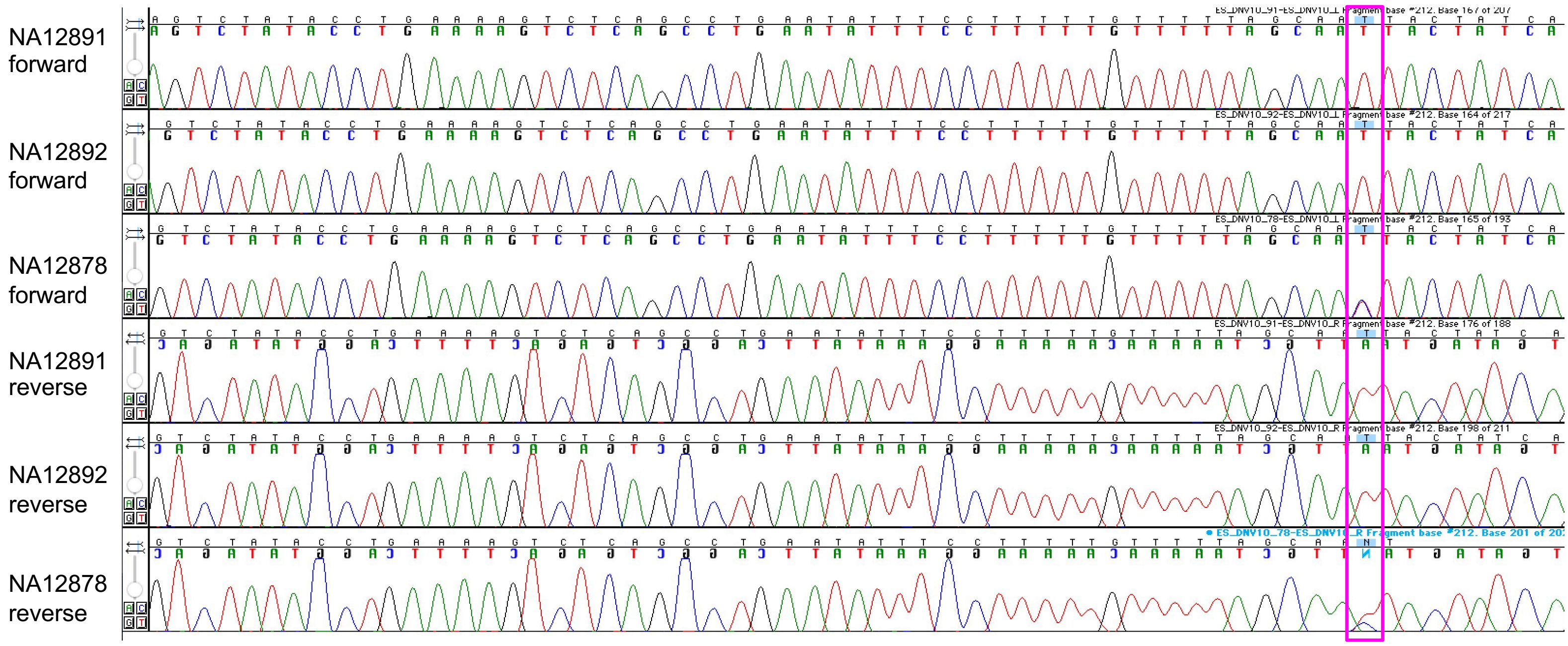

Figure S11

chr4_65406505_A_G

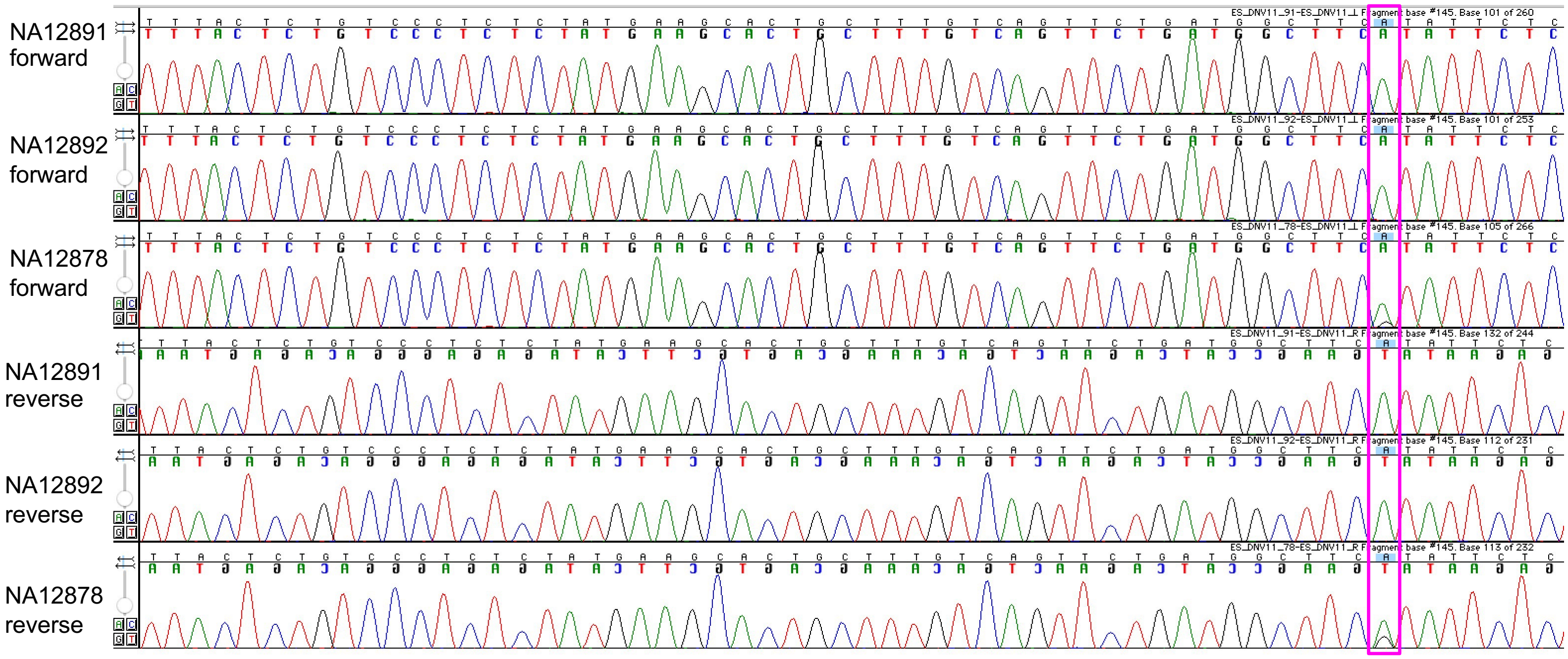

Figure S12

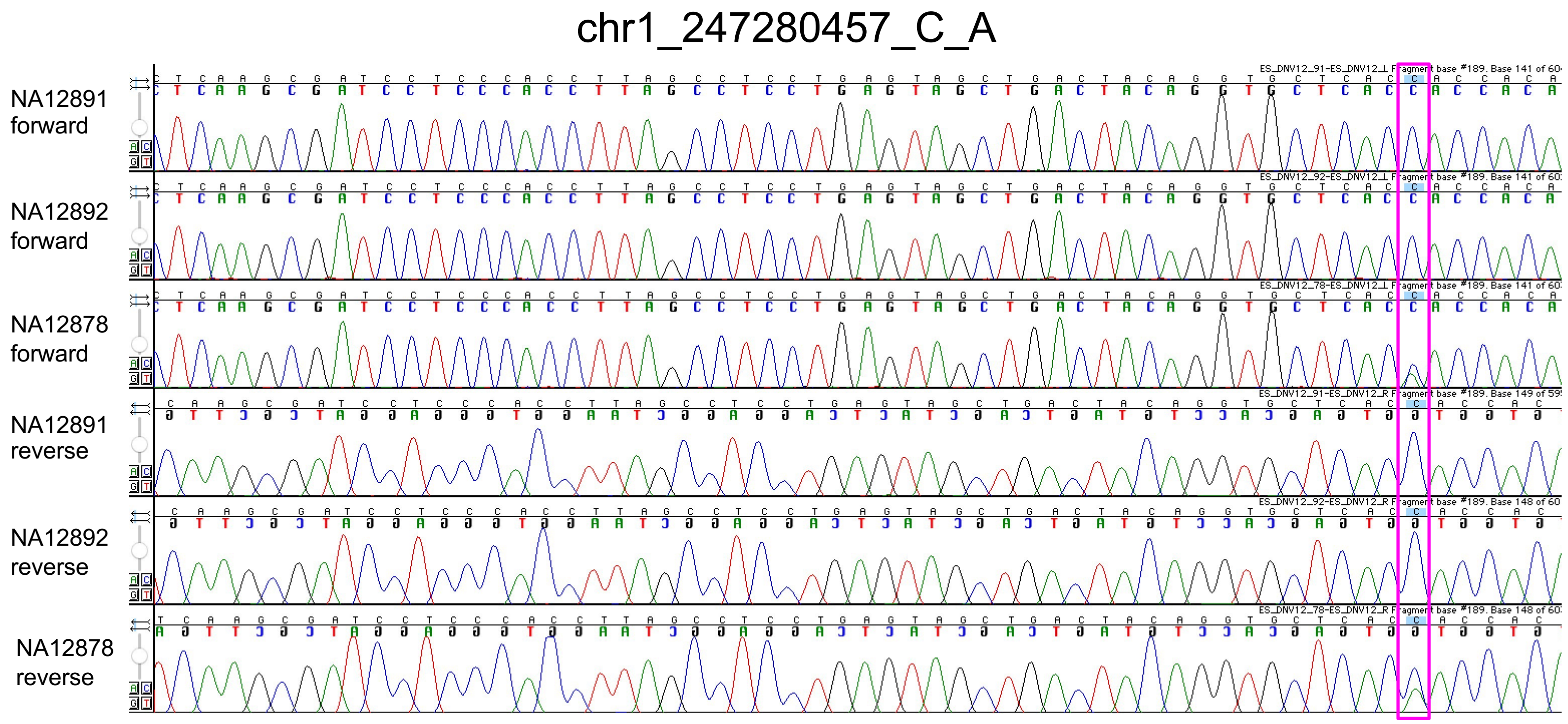
chr1_247280457_C_A
NA12891 forward
NA12892 forward
NA12878 forward
NA12891 reverse
NA12892 reverse
NA12878 reverse

### Figure S13

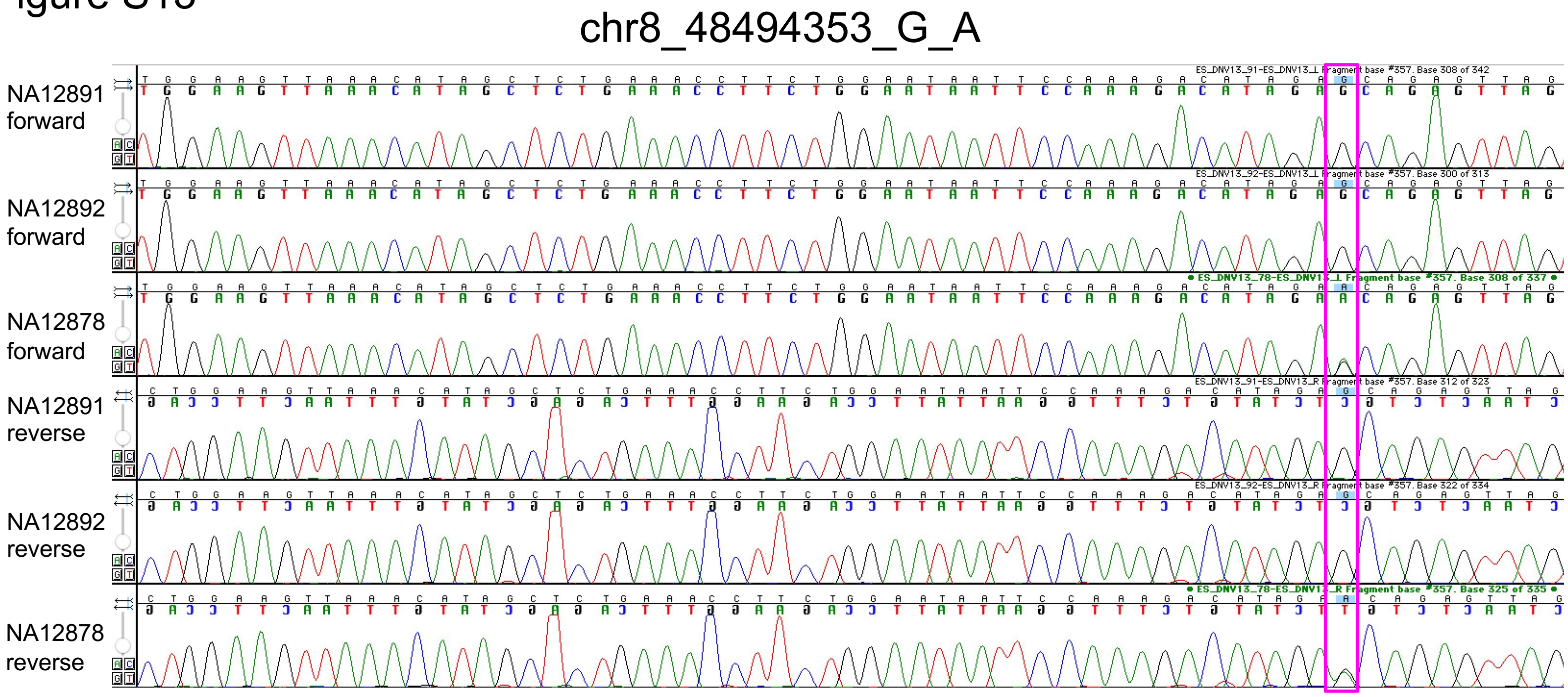
chr8_48494353_G_A
NA12891 forward
NA12892 forward
NA12878 forward
NA12891 reverse
NA12892 reverse
NA12878 reverse

Figure S14

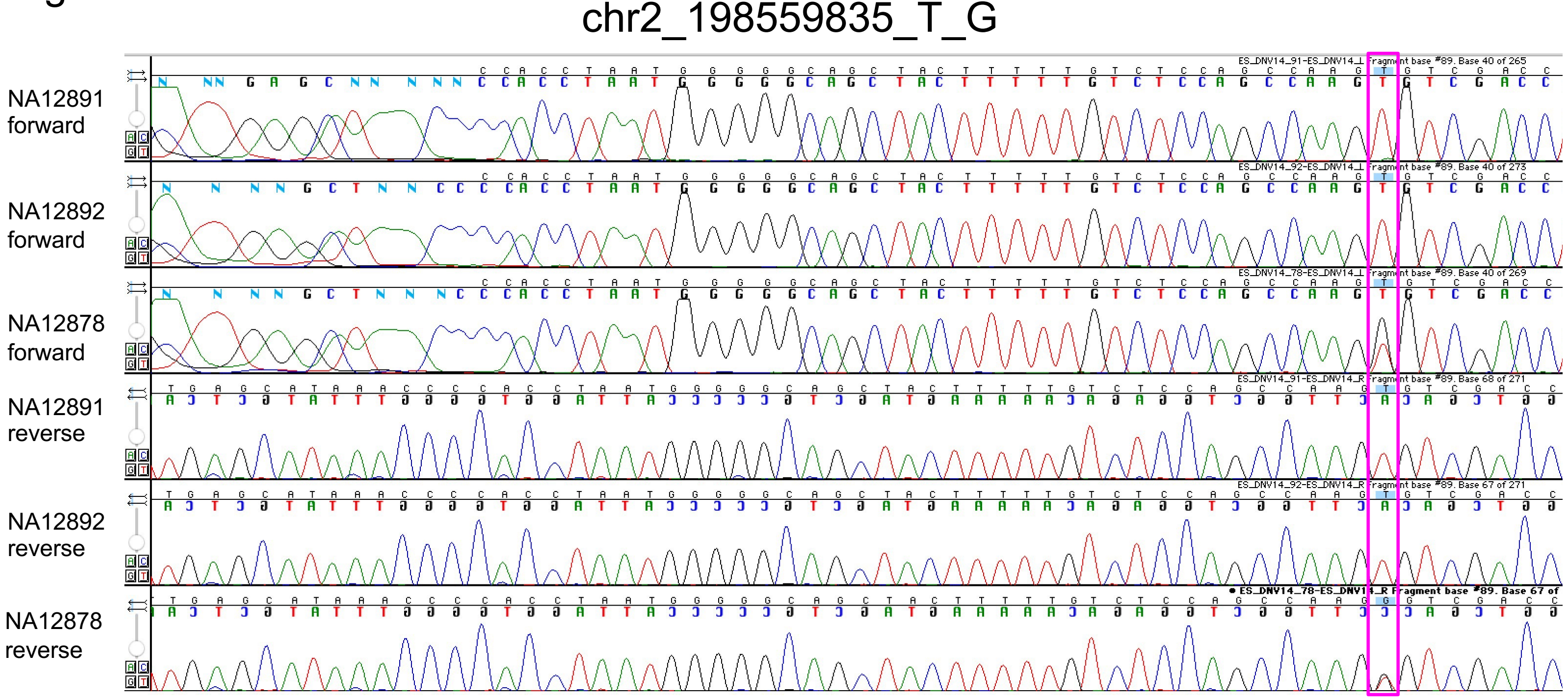
chr2_198559835_T_G
NA12891 forward
NA12892 forward
NA12878 forward
NA12891 reverse
NA12892 reverse
NA12878 reverse

Figure S15

#### chr4_5461303_G_A

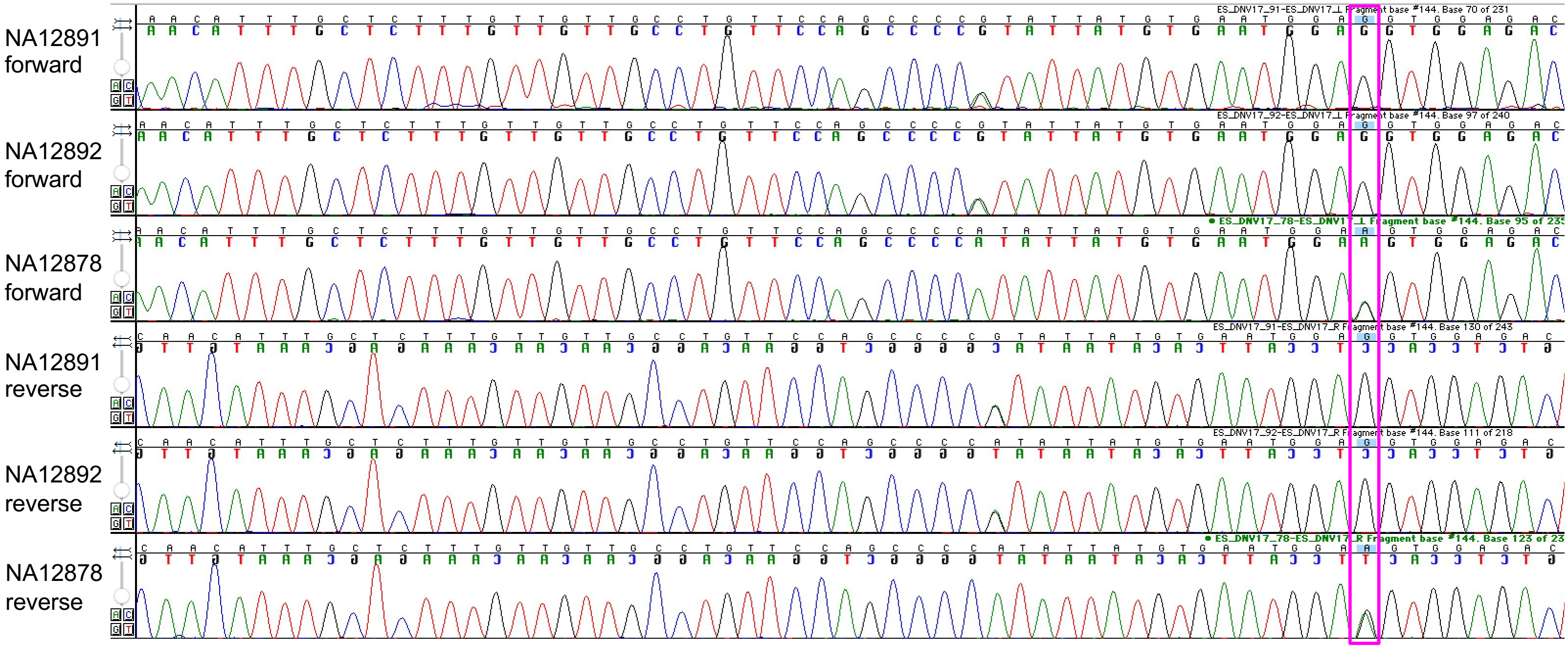

Figure S16

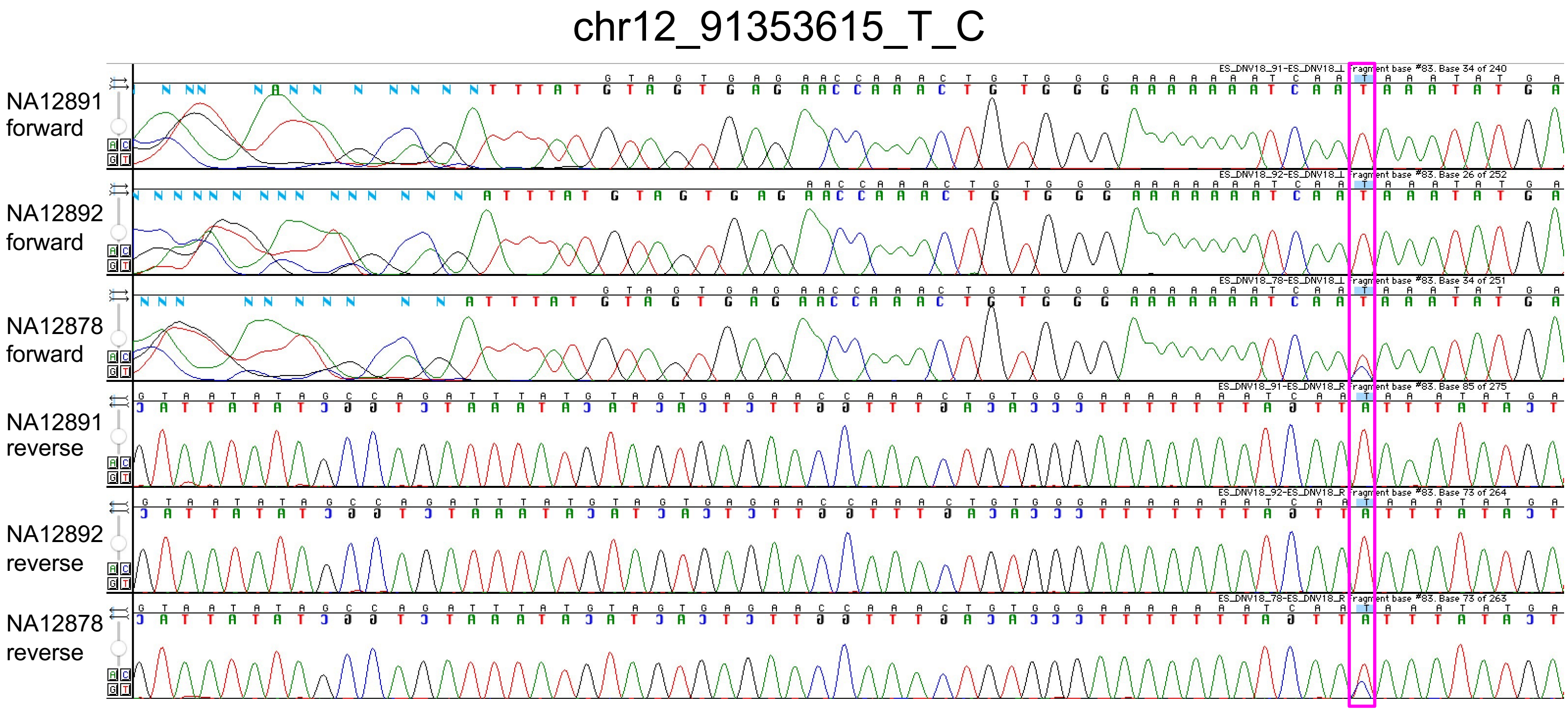

chr12_91353615_T_C
NA12891 forward
NA12892 forward
NA12878 forward
NA12891 reverse
NA12892 reverse
NA12878 reverse

### Figure S17

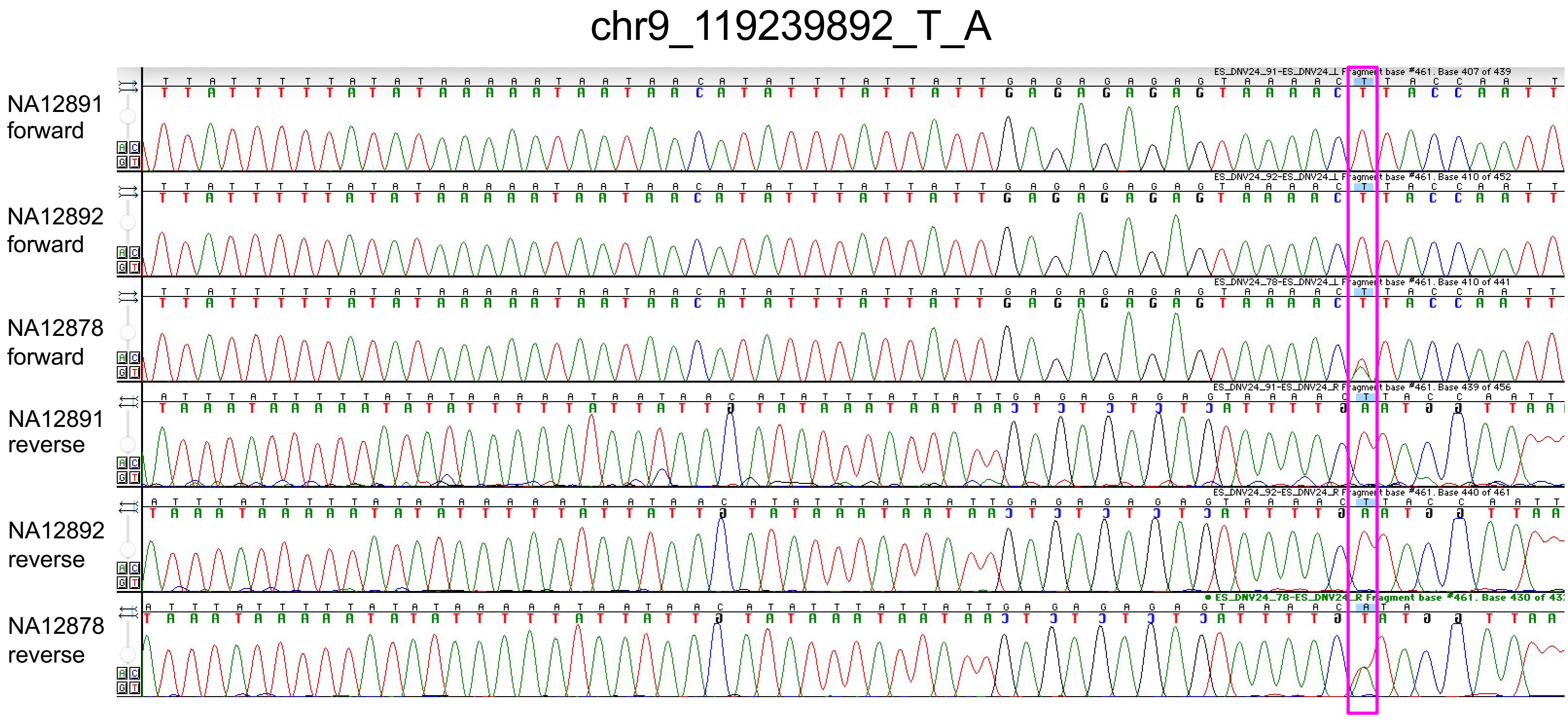
chr9_119239892_T_A
NA12891 forward
NA12892 forward
NA12878 forward
NA12891 reverse
NA12892 reverse
NA12878 reverse

Figure S18

chr10_109204409_C_T

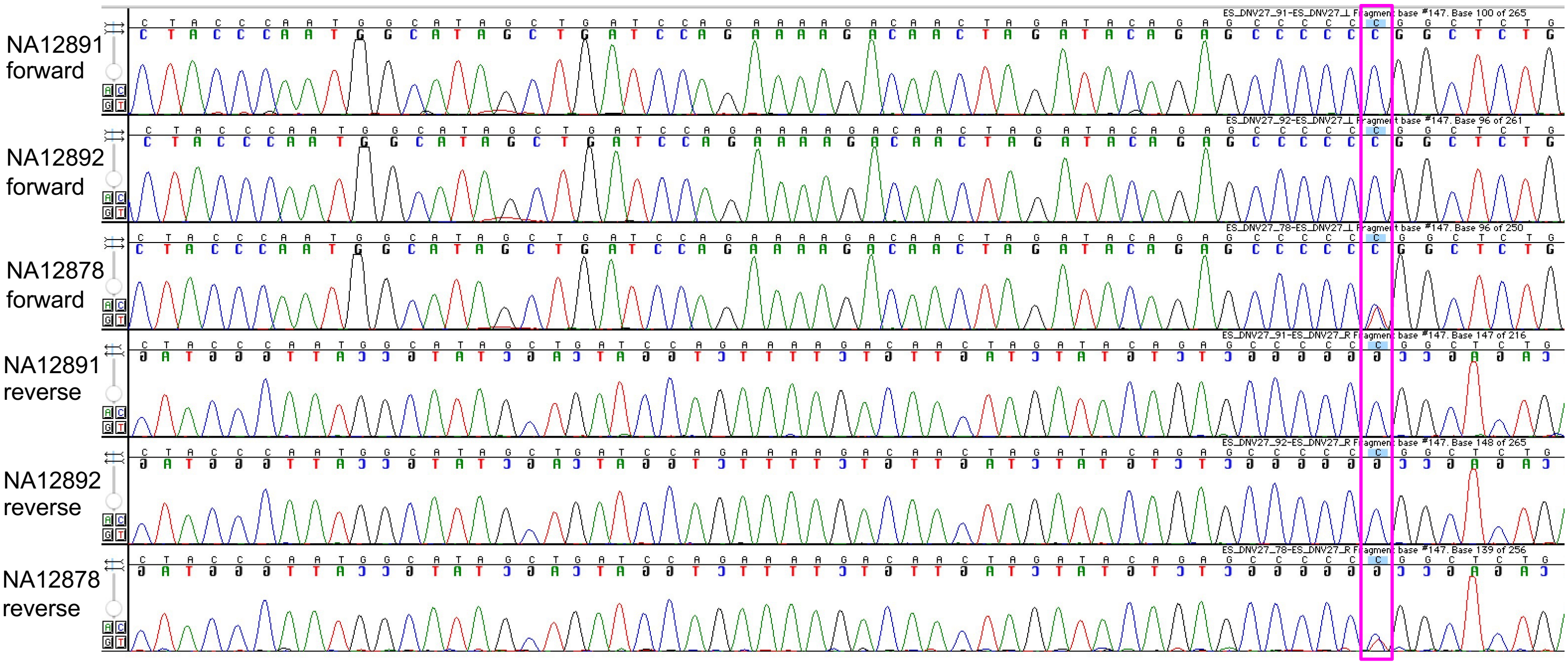

### Figure S19

#### chr6_71729112_T_C

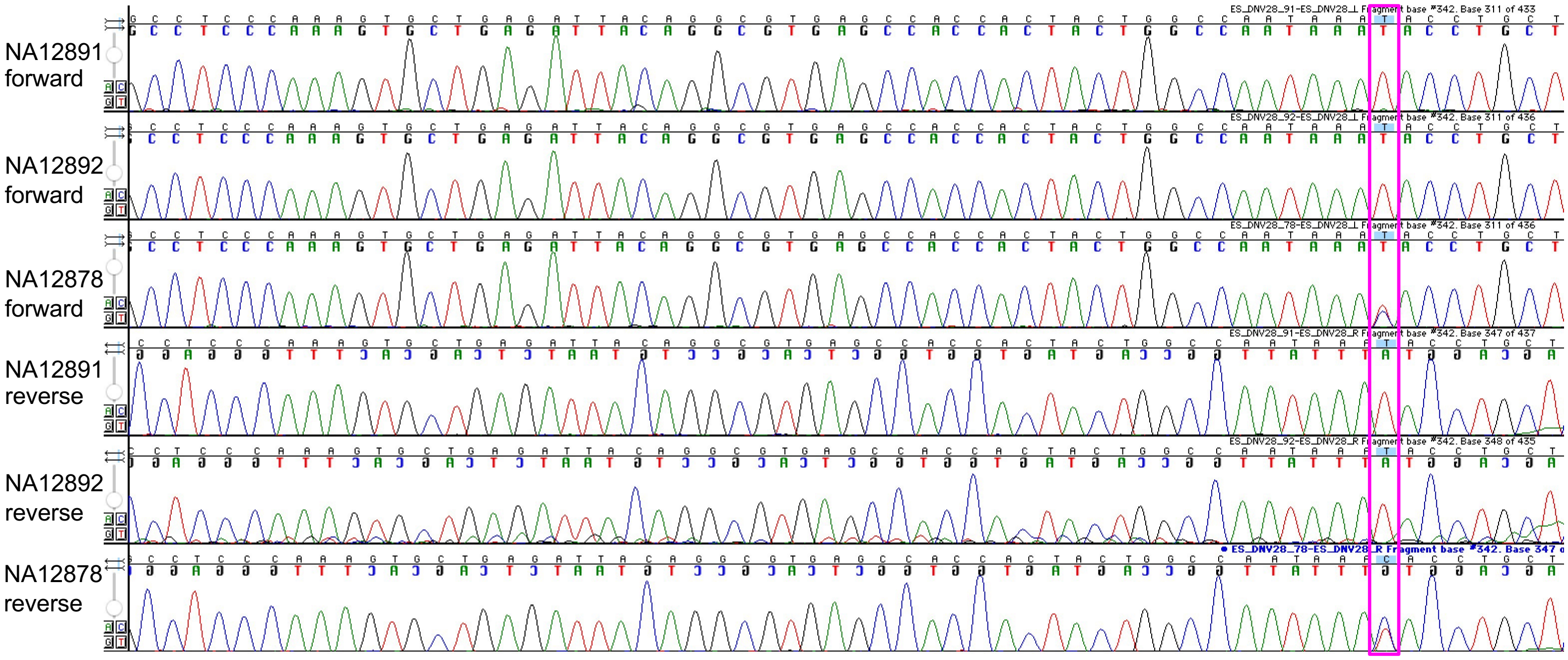

### Figure S20

chr13_81142986_A_G

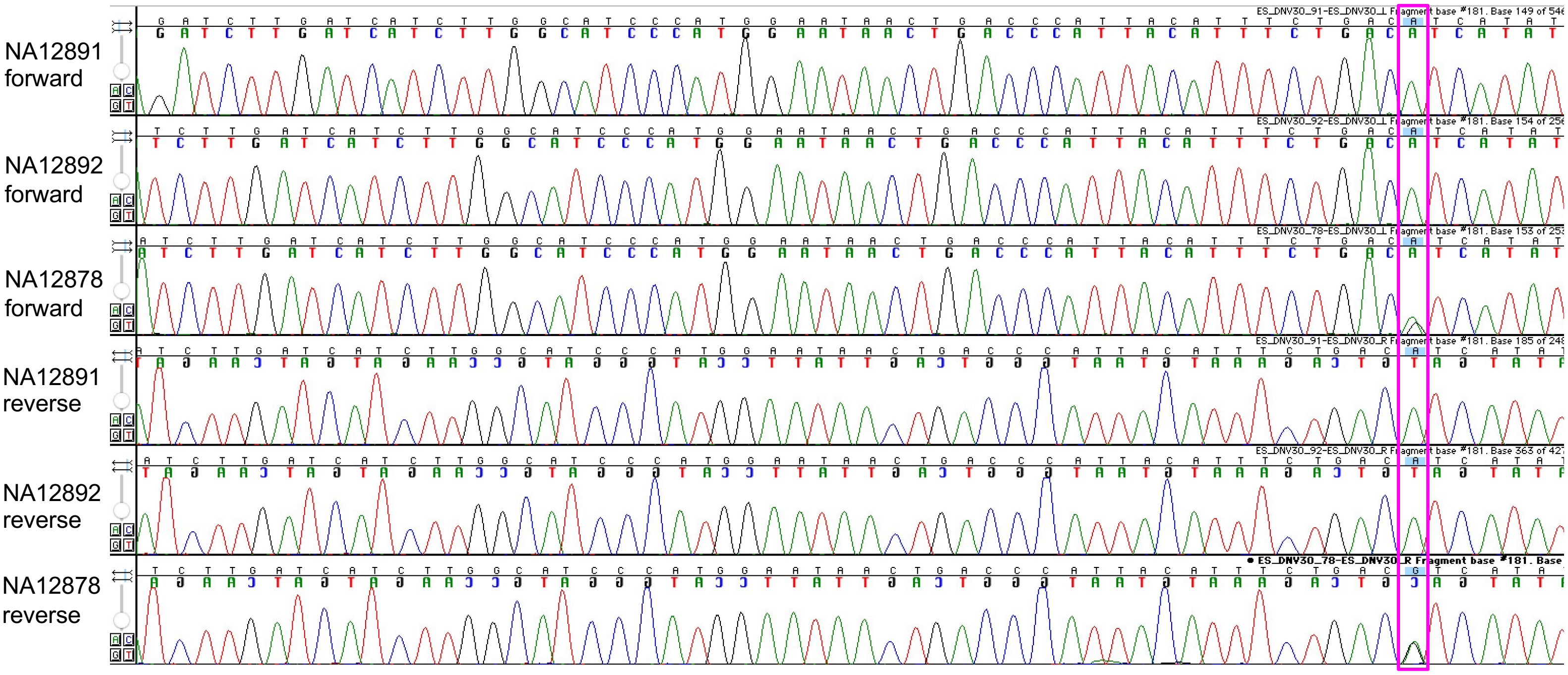

### Figure S21

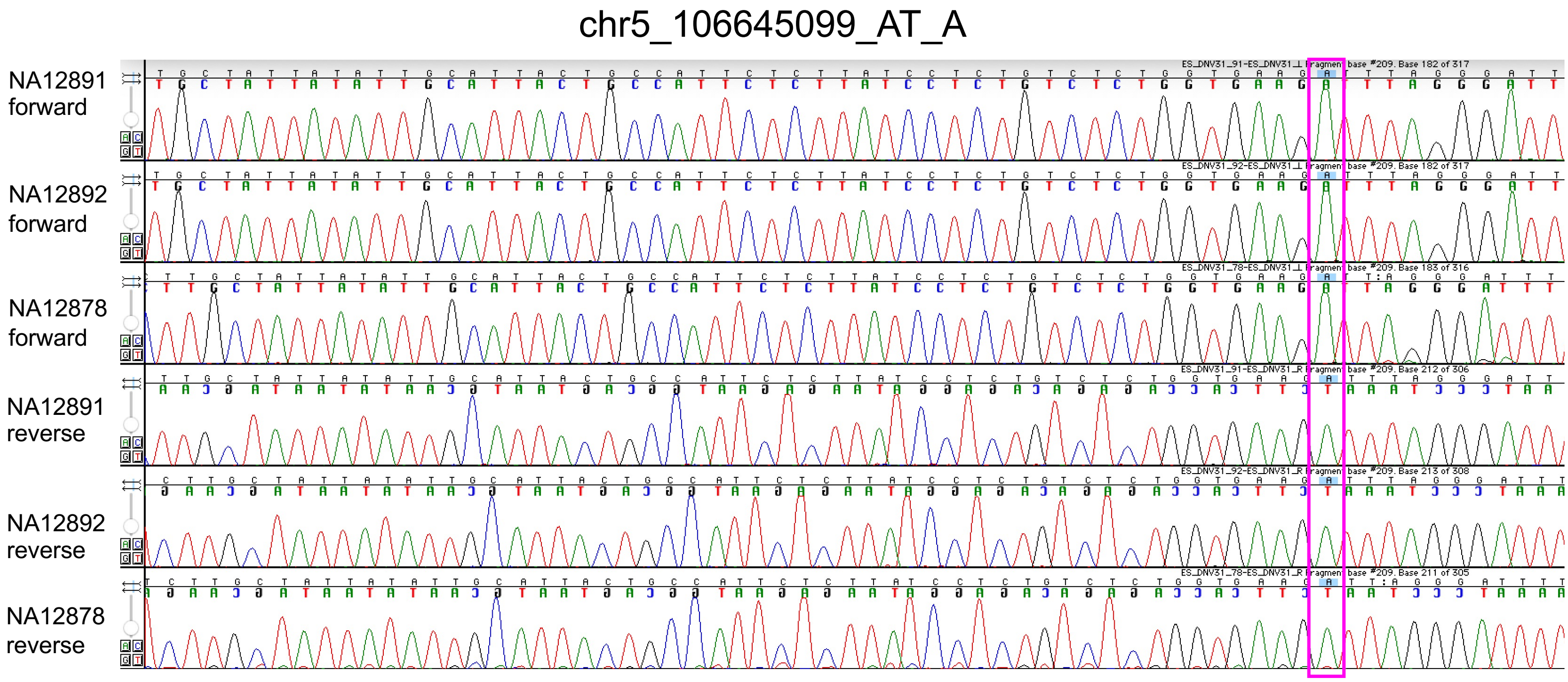
chr5_106645099_AT_A
NA12891 forward
NA12892 forward
NA12878 forward
NA12891 reverse
NA12892 reverse
NA12878 reverse

### Figure S22

#### chr5_123998792_A_C

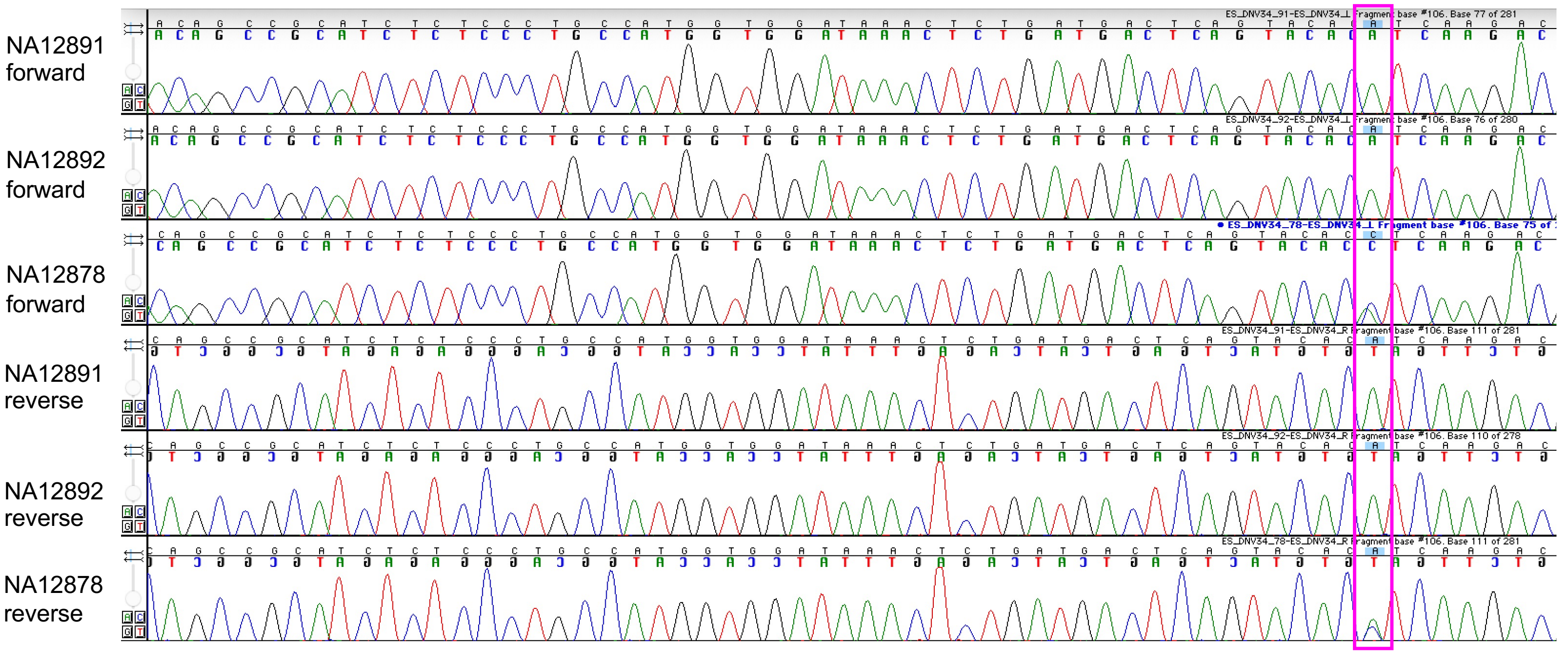

### Figure S23

chr19_36641978_C_T

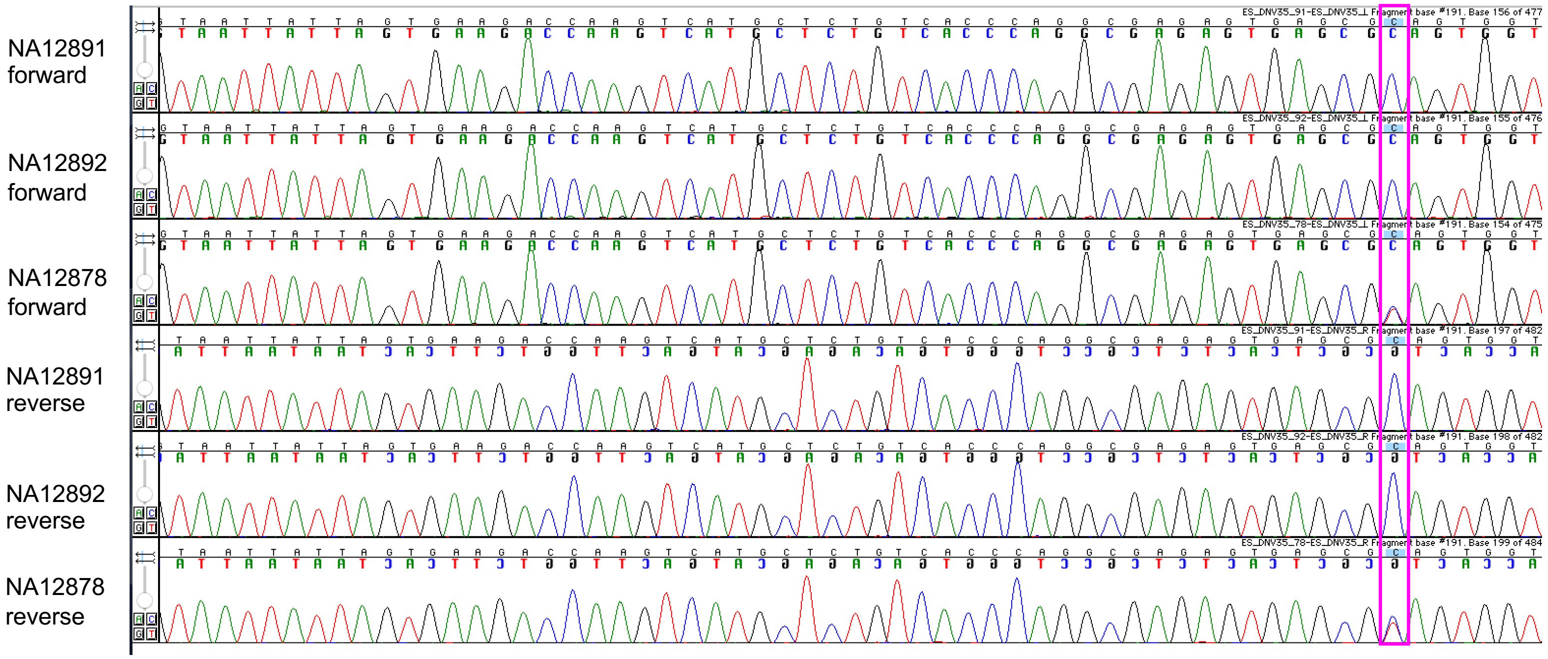

### Figure S24

#### chr13_71846473_C_A

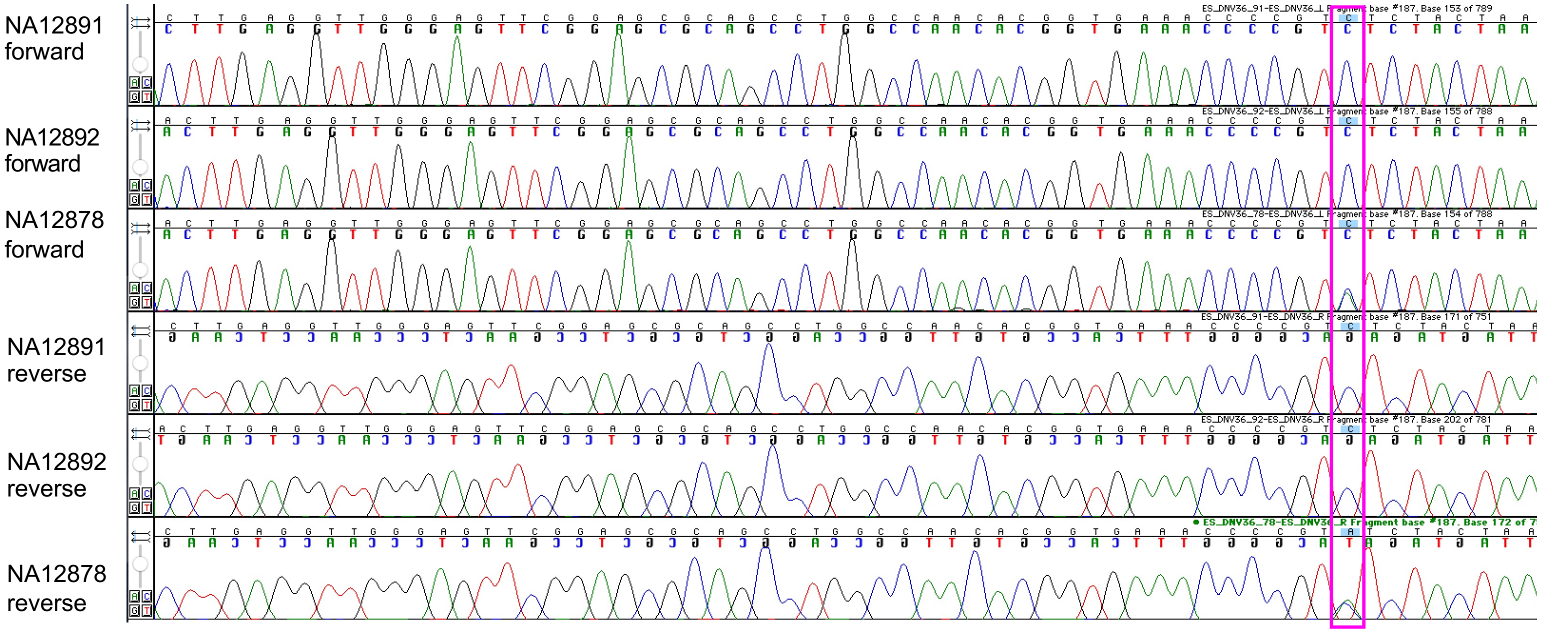

### Figure S25

#### chr2_155841078_G_A

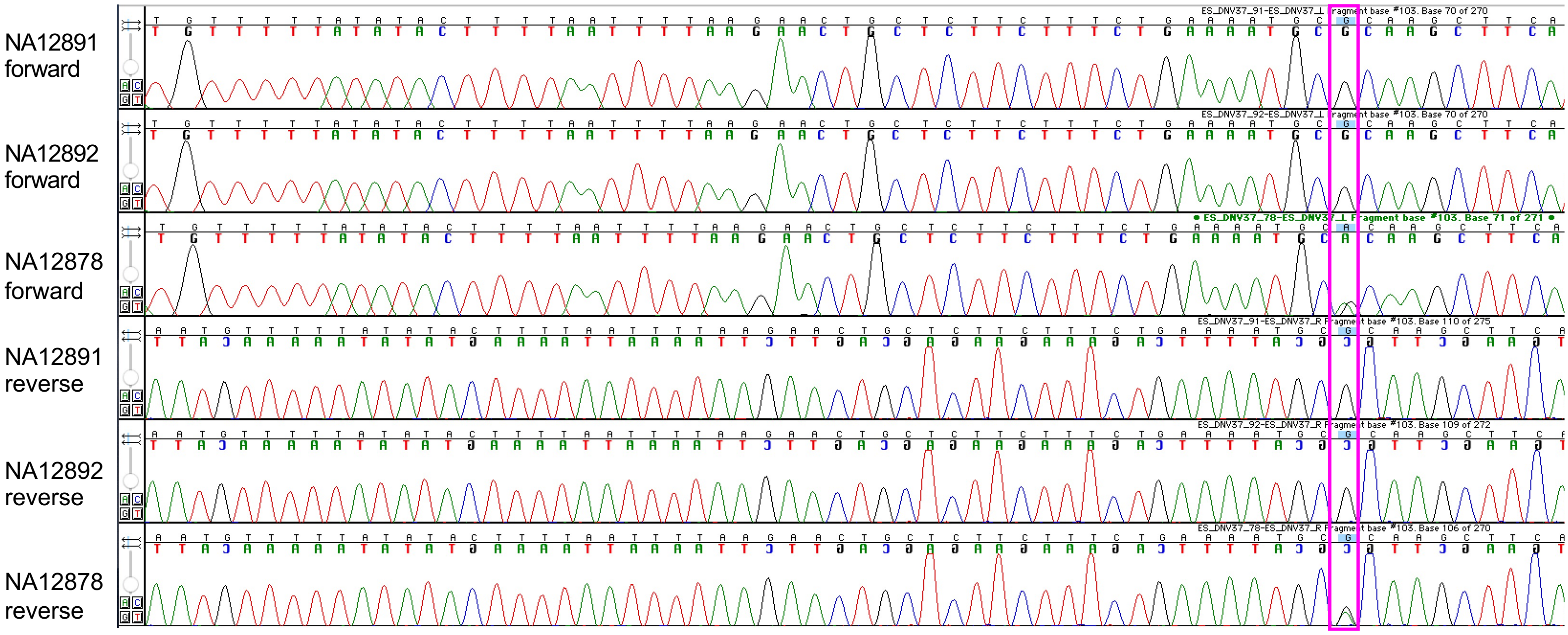

### Figure S26

#### chr20_24046519_G_A

NA12891 forward

NA12892 forward

NA12878 forward

NA12891 reverse

NA12892 reverse

NA12878 reverse

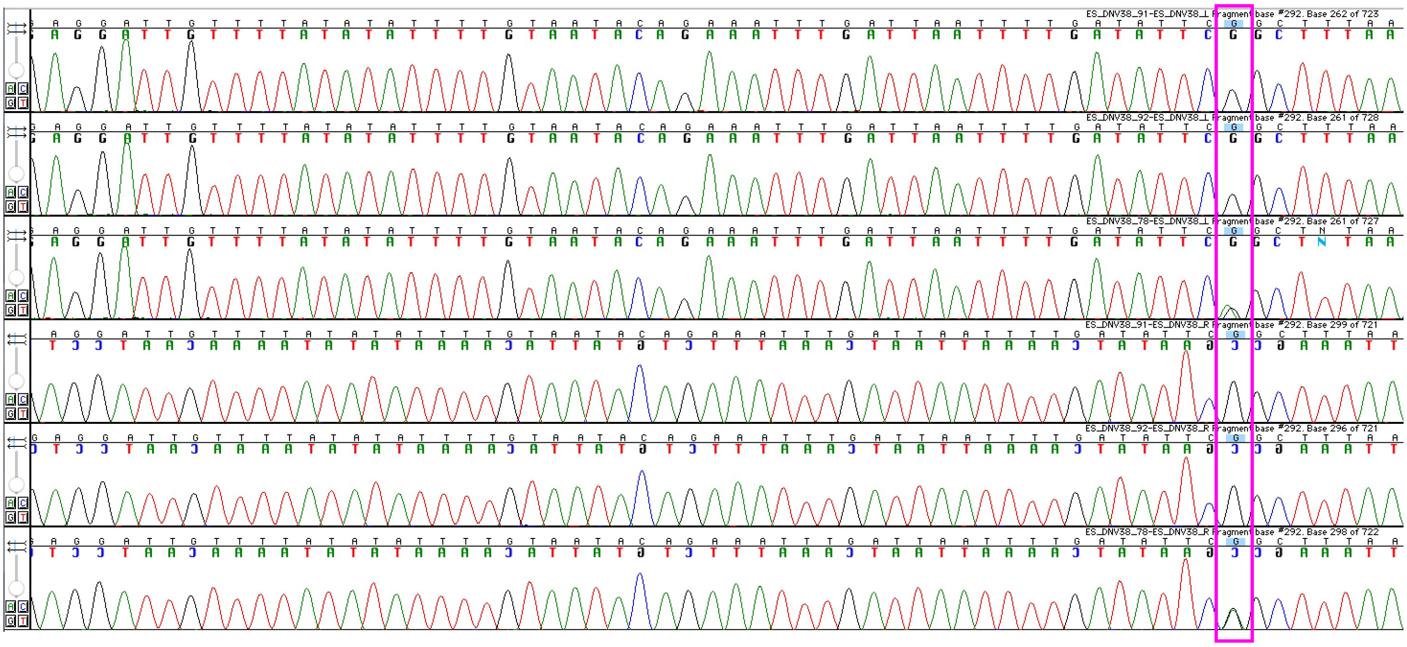

### Figure S27

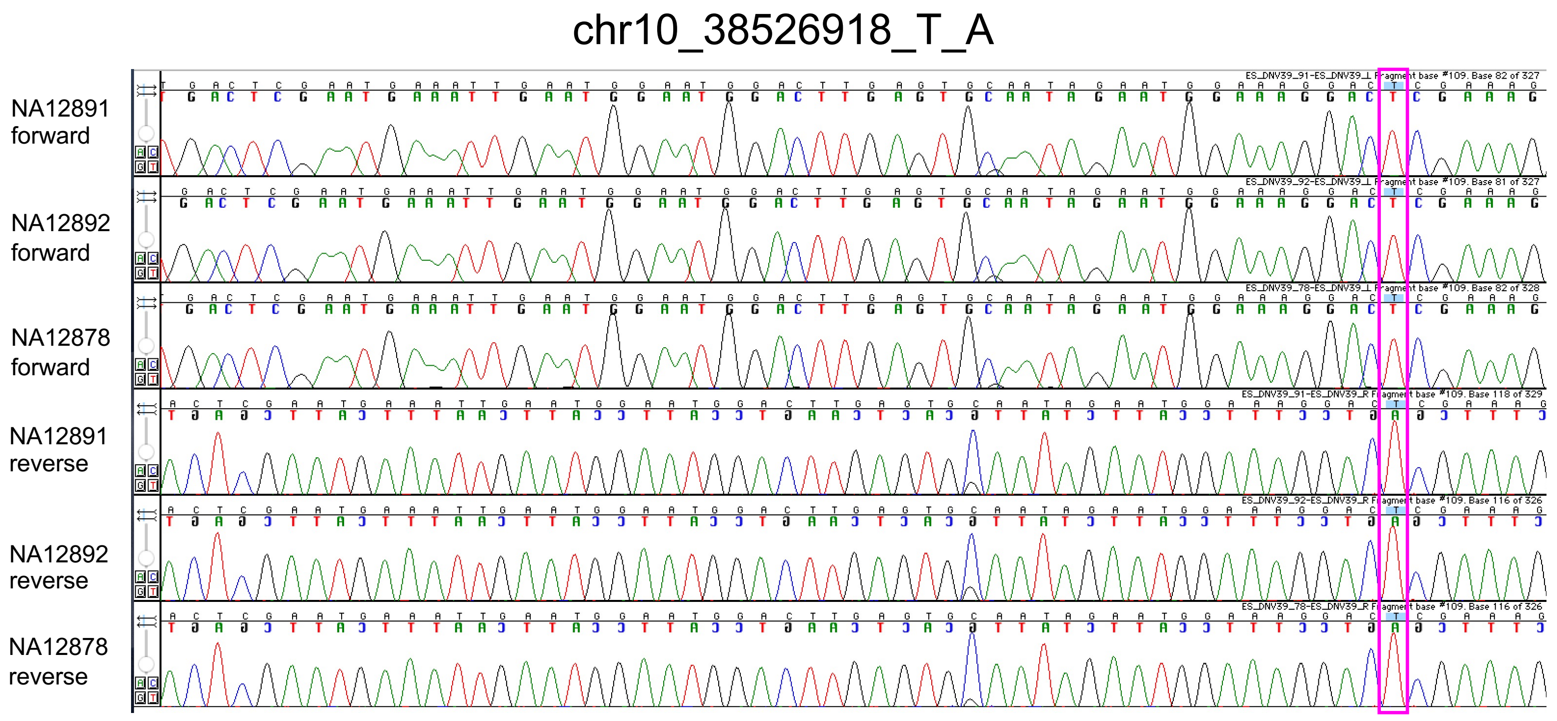


Note: This variant is not real

### Figure S28

chr11_134531608_C_G

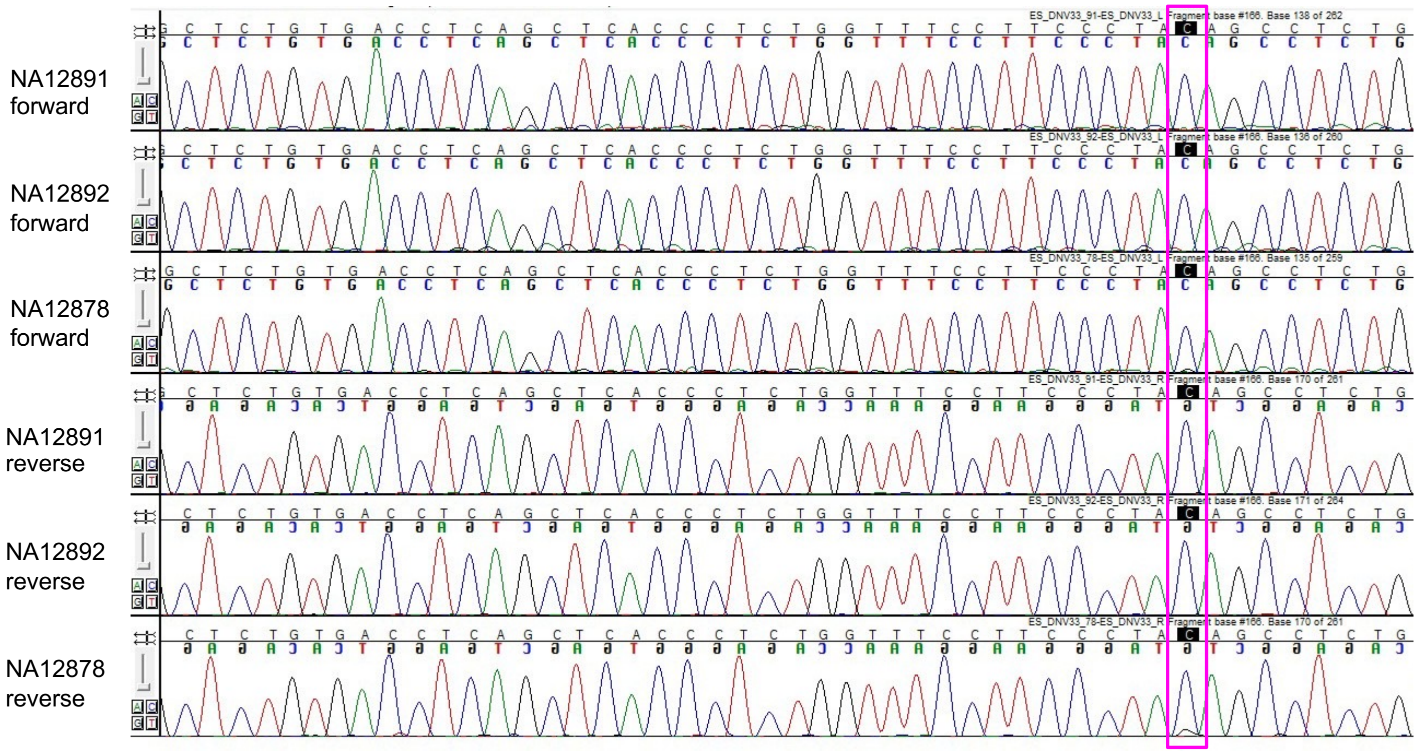


Note: Sanger results inconclusive, see our follow up ONT sequencing results

Figure S29

chr21_19977497_C_T

NA12891 forward

NA12892 forward

NA12878 forward

NA12891 reverse

NA12878 reverse

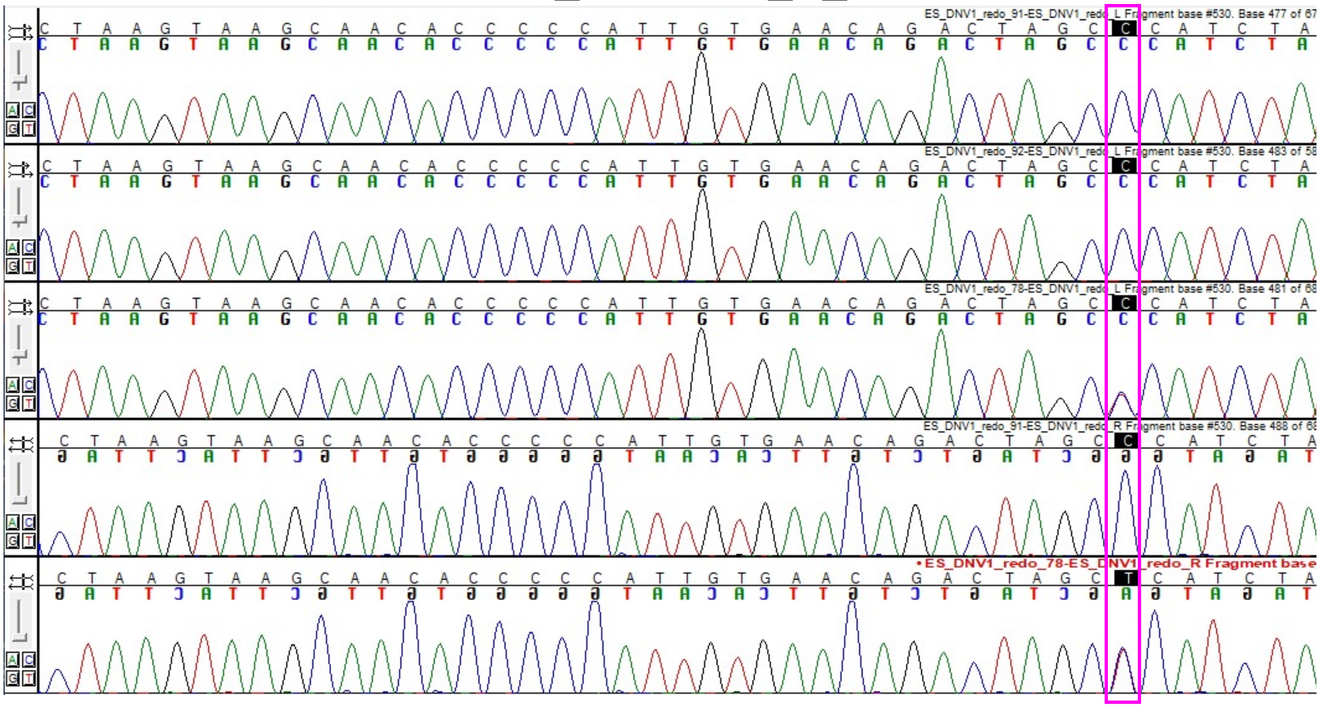

Note: No data for NA12892 reverse

Figure S30

IGV images of Oxford Nanopore Technologies (ONT) sequencing of amplicon

chr11_134531608_C_G

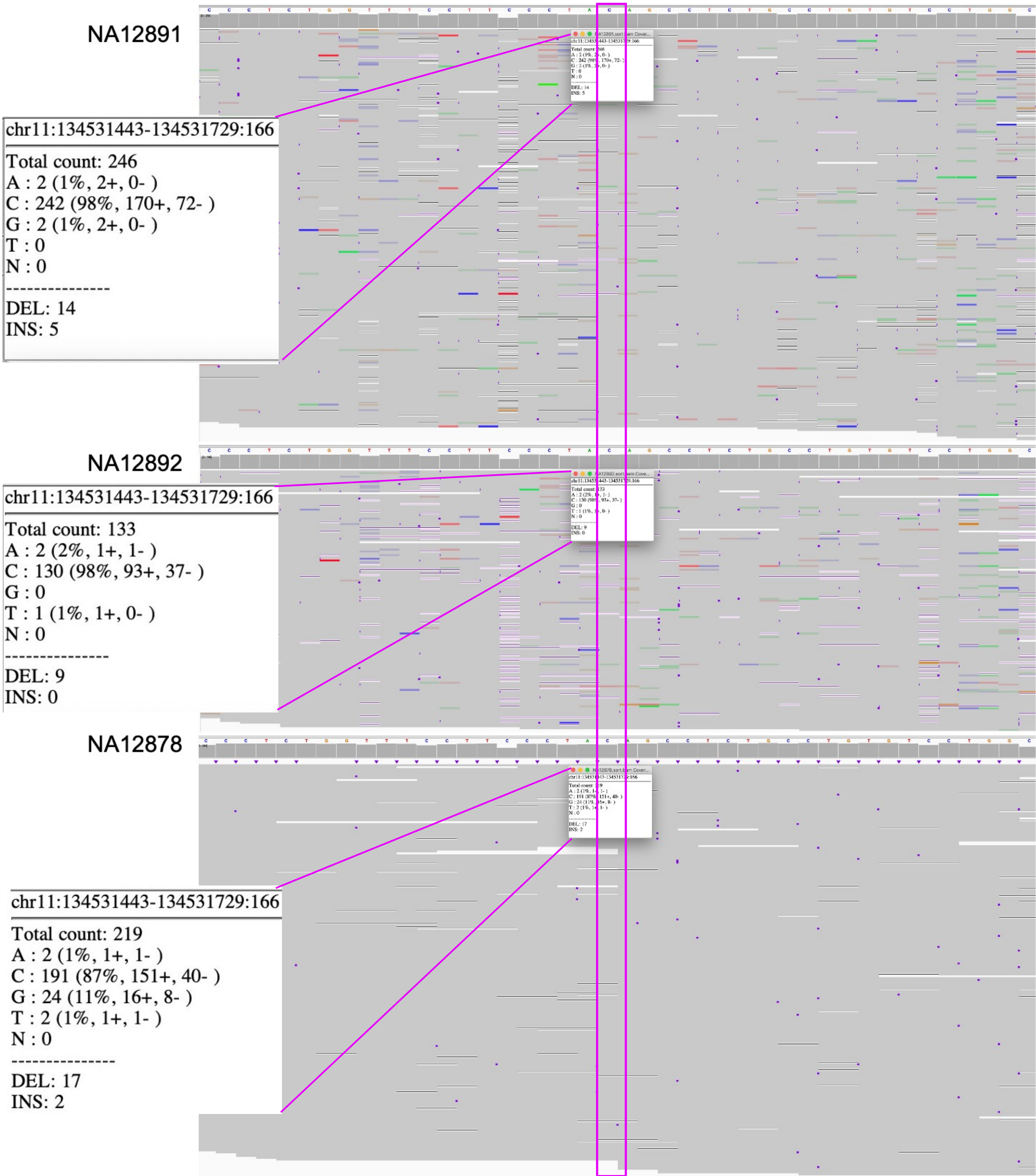

### Figure S31

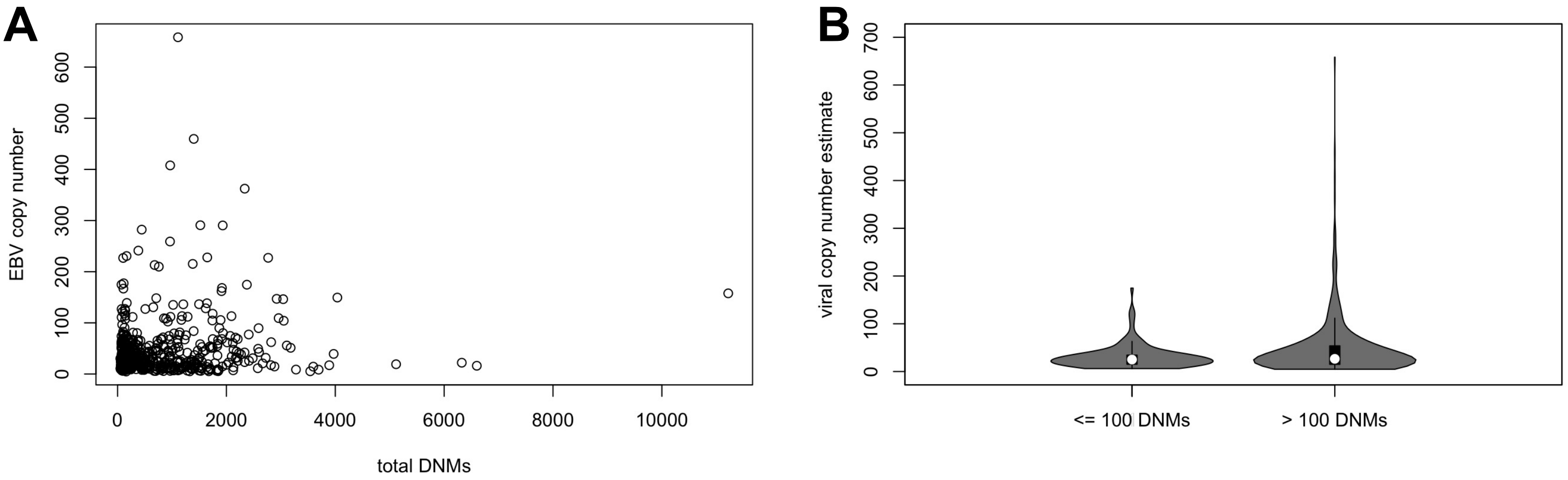
A
EBV copy number
0
100
200
300
400
500
600
0
2000
4000
6000
8000
10000
total DNMs
B
viral copy number estimate
0
100
200
300
400
500
600
700
<= 100 DNMs
> 100 DNMs

Figure S32

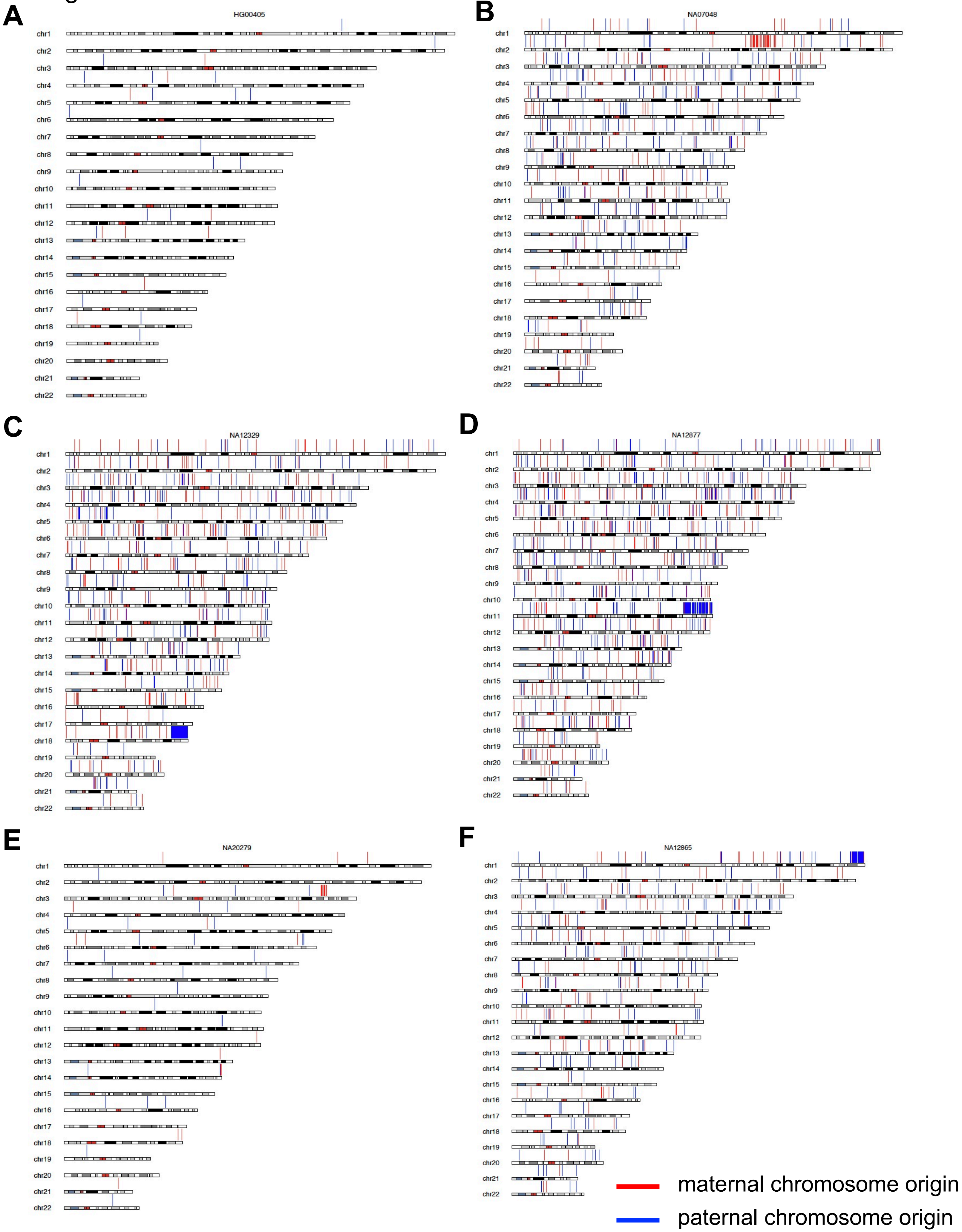
A
HG00405
B
NA07048
C
NA12329
D
NA12877
E
NA20279
F
NA12865
chr1
chr2
chr3
chr4
chr5
chr6
chr7
chr8
chr9
chr10
chr11
chr12
chr13
chr14
chr15
chr16
chr17
chr18
chr19
chr20
chr21
chr22
maternal chromosome origin
paternal chromosome origin

Figure S33

A

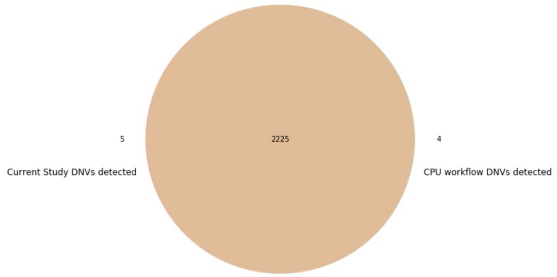


B

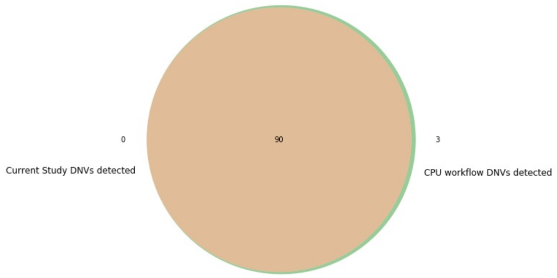


C

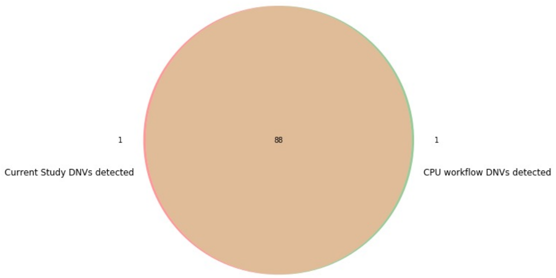

Figure S34

A

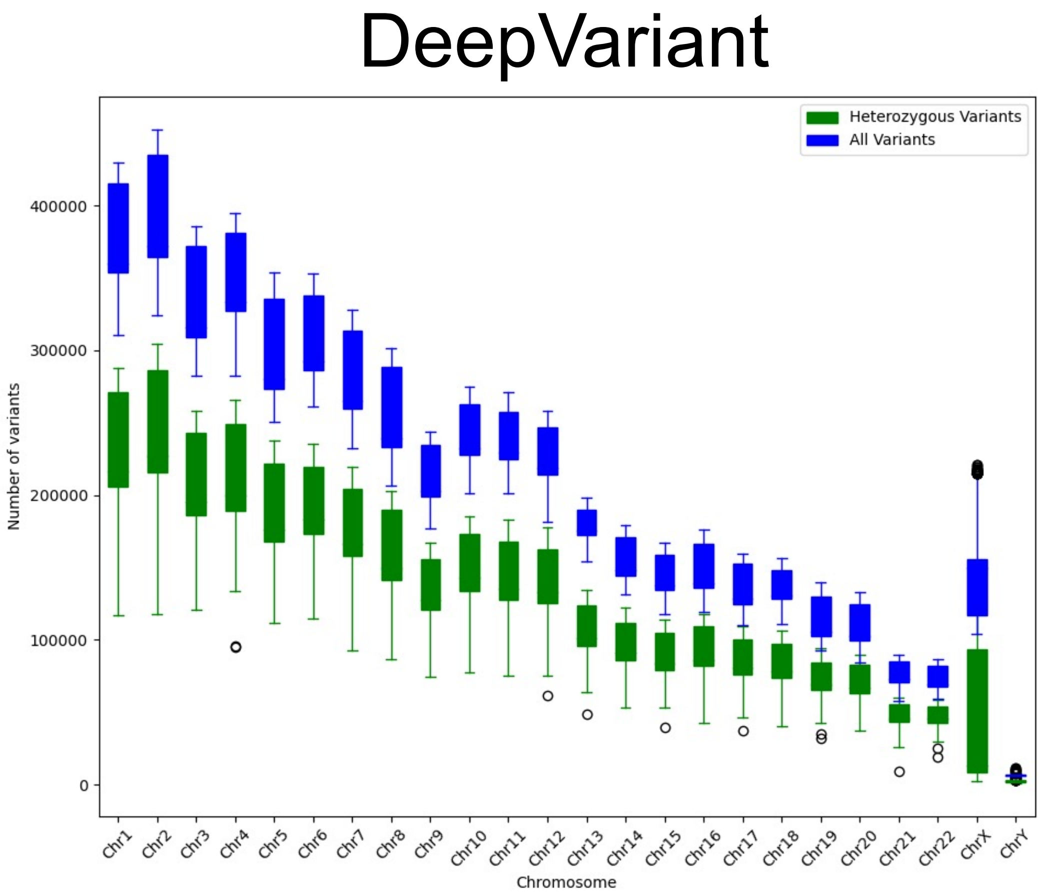


B

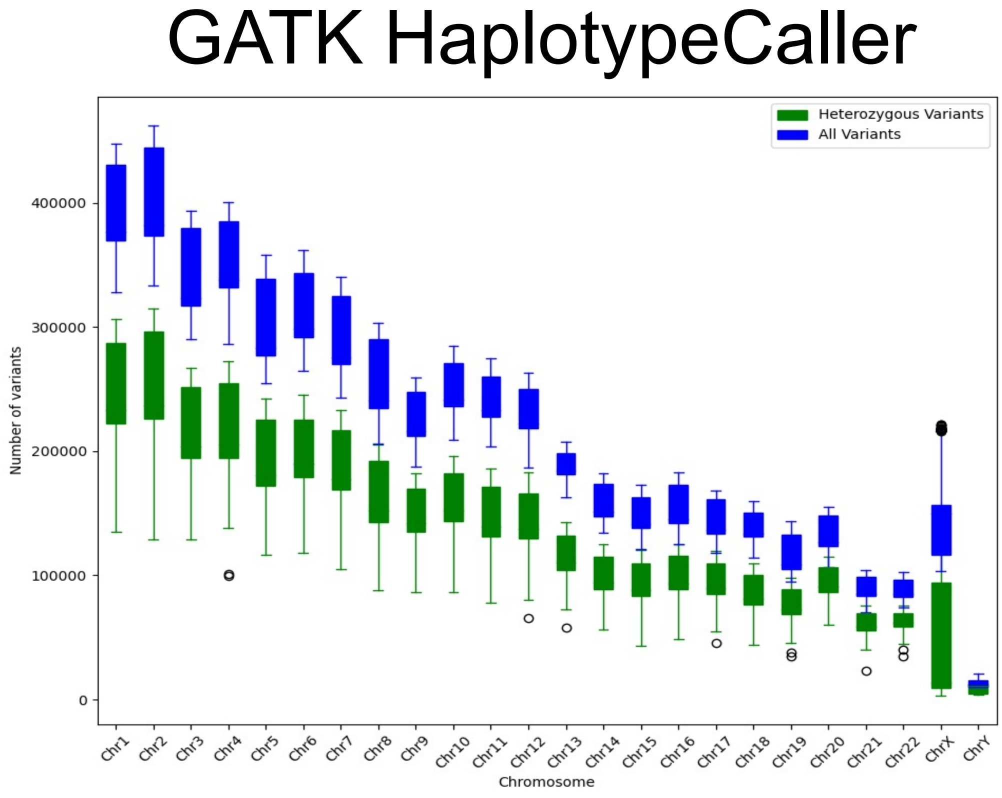

Figure S35

DeepVariant

**A**

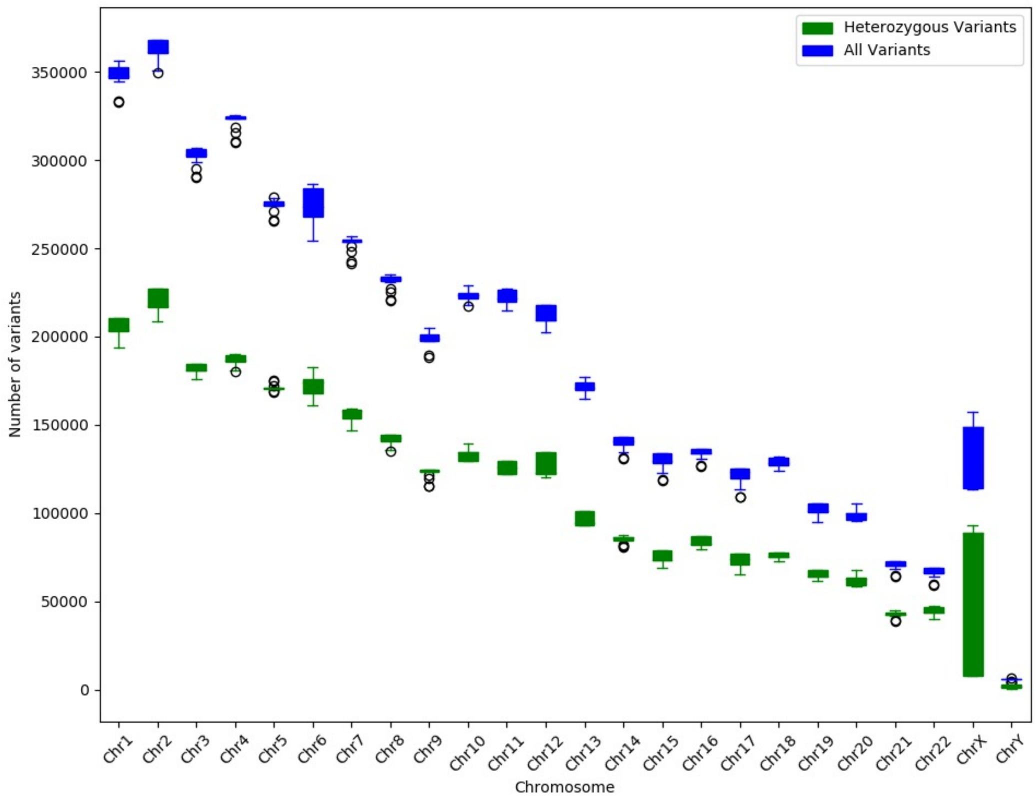


GATK HaplotypeCaller

**B**

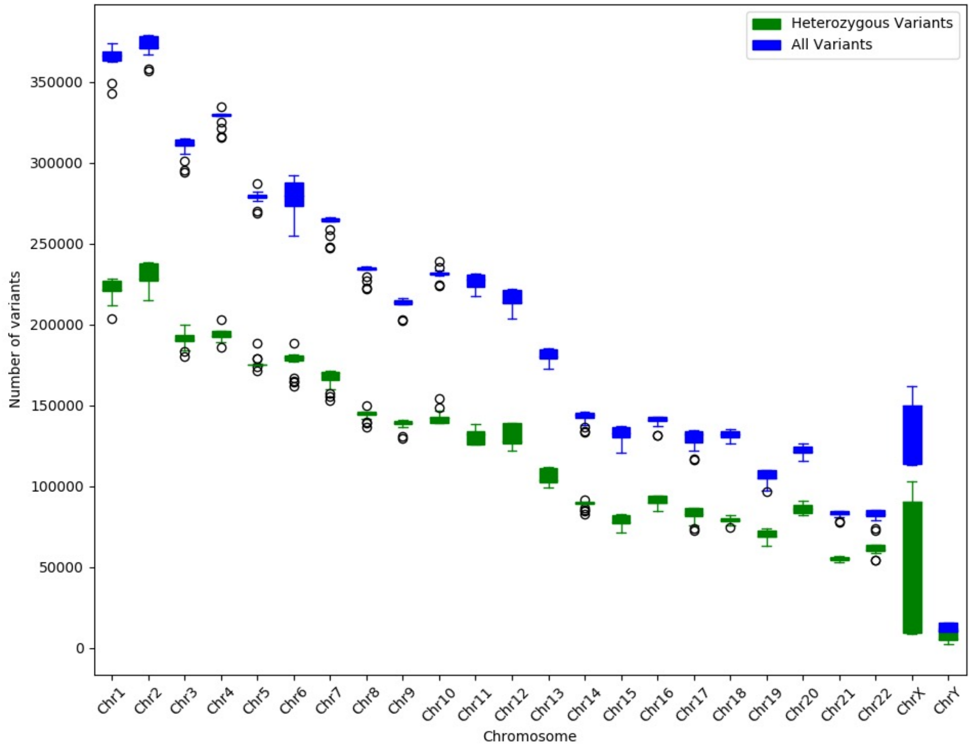

### Figure S36

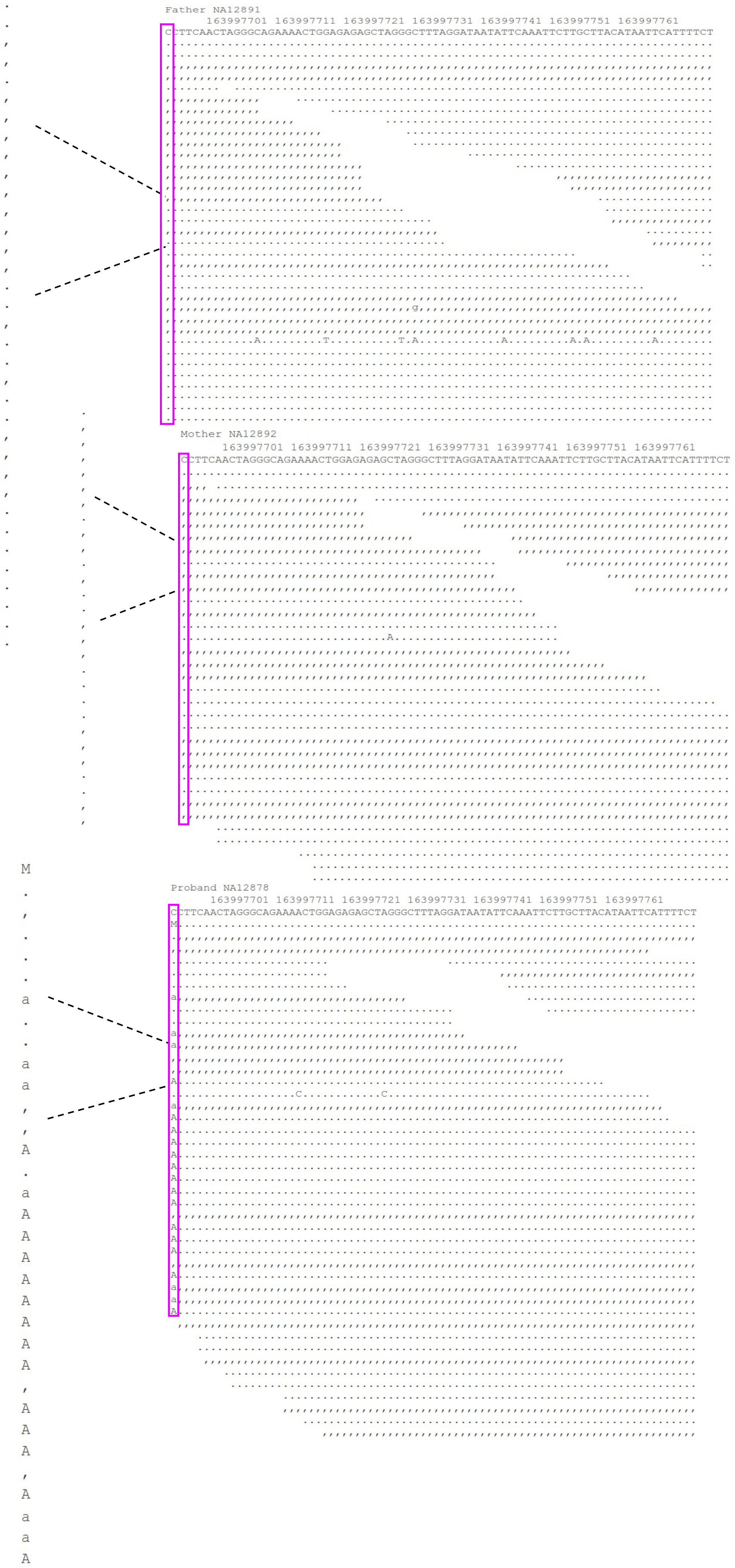

Father NA12891
163997701 163997711 163997721 163997731 163997741 163997751 163997761
CCTTCAACTAGGGCAGAAAACTGGAGAGAGCTAGGGCTTTAGGATAATATTCAAATTCTTGCTTACATAATTCATTTTCT
Mother NA12892
163997701 163997711 163997721 163997731 163997741 163997751 163997761
CCTTCAACTAGGGCAGAAAACTGGAGAGAGCTAGGGCTTTAGGATAATATTCAAATTCTTGCTTACATAATTCATTTTCT
Proband NA12878
163997701 163997711 163997721 163997731 163997741 163997751 163997761
CCTTCAACTAGGGCAGAAAACTGGAGAGAGCTAGGGCTTTAGGATAATATTCAAATTCTTGCTTACATAATTCATTTTCT
