## Supplemental Table Legends for "*de novo* variant calling identifies cancer mutation profiles in the 1000 Genomes Project"

**Supplement Table Legends**

**S1 DNV scoring by visual inspection of underlying reads with SAMtools tview**
DNVs from four 1000G trios were scored by visual assessment of read data at each variant position in the father, mother, and child.

**S2 Information about the 602 trios for 1000G**
Shown are the family identifiers, individual identifiers, relationship to other family members, sex, population and superpopulation for each individual in the 602 trios.

**S3 DNV callset in 1000G**
The file name and md5 for the Table S3 are shown in the excel workbook. This a reference to the actual file that is separate from the workbook due to size.

**S4 Total DNVs per individual in 1000G**
Shown are the counts of DNVs per individual in the 1000G as well as the category to which they belong in terms of counts.

**S5 Total DNVs per individual in SSC**
Shown are the counts of DNVs per individual in the SSC.

**S6 Phasing results for the 1000G**
The file name and md5 for the Table S5 are shown in the excel workbook. This a reference to the actual file that is separate from the workbook due to size.

**S7 Results of Sanger and ONT sequencing in NA12878**
The twenty-five variants chosen for assessment by Sanger sequencing are shown in this table. Additionally information is present of the one variant also sequenced by ONT technology.

**S8 Mutation profile signatures in 1000G**
The results of mutation profile analysis for each child in 1000G is shown.

**S9 Read-depth based karyotype results for 1000G**
For each individual, the predicted karyotype based on read-depth is shown.

**S10 DNVs in DNA damage genes**
DNVs found in individuals from 1000G occurring in DNA damage genes.

**S11 Read-depth based estimateion of EBV copy number**
For each individual, the predicted EBV copy number based on read-depth is shown.

**S12 Results of the two-phase DNV per gene enrichment analysis in 1000G**
Shown are the results of running the chimpanzee-human and denovolyzeR tests on DNVs identified in 1000G (phase 1). Each p-value shown is the Bonferroni corrected value for multiple tests. Also, shown is the result of phase 2 that compared individuals

with <= 100 DNVs to those with > 100 DNVs. Only one gene was significant in both phases and that was *IGLL5*.

**S13 Protein-coding DNVs in *IGLL5***

Shown are the *IGLL5* protein-coding DNVs identified in individauls from 1000G.

**S14 Clinical DNVs**

Shown are the DNVs meeting clinical relevance. These DNVs are important to consider when using the 1000G as a resource to filter variants in clinic.
