## Supplemental Figure Legends for "*de novo* variant calling identifies cancer mutation profiles in the 1000 Genomes Project"

### Supplement Figures Legends

**S1 Sample population distribution**
A distribution of the populations by super and sub populations defined by the 1000 Genomes Project (1000G). Blue represents Ad Mixed American super population, red represents African super population, purple is East Asian super population, grey represents European super population, and green represents South Asian super population.

**S2 Phase distribution of DMV in the 1000G**

A) Histogram of the percent autosomal DMVs that was fixed
B) A distribution of percent autosomal DMVs phased vs. total autosomal DMVs.
C) Violin plot of percent of autosomal DMVs that with paternal parent-of-origin.
D) Distribution of DMVs with paternal parent-of-origin. The pink graph represents samples that had greater than 100 DMVs. The blue graph represents samples that had less than or equal to 100 DMVs. There is a trend of higher percent of DMVs with paternal parent-of-origin compared in the group that had less than or equal 100 DMVs compared to those with greater than 100 DMVs, which would be expected if the DMVs are real.

**S3 Phase distribution of DMV in Simons Simplex Collection**

A) Histogram of the percent autosomal DMVs that was fixed in the Simons Simplex Collection (SSC).
B) A distribution of percent autosomal DMVs phased vs. total autosomal DMVs.
C) Violin plot of percent of autosomal DMVs that with paternal parent-of-origin.
D) Distribution of DMVs with paternal parent-of-origin. The pink graph represents samples that had greater than 100 DMVs. The blue graph represents samples that had less than or equal to 100 DMVs.

**S4 Comparison of distribution of allele balance per trio between 1000 Genomes Project and SSC**
Distribution of the allele balance for each DNV between the 602 trios of the 1000G, in blue, and the 4216 of the SSC in red. The allele balance is closer to 0.5 for SSC compared to 1000G.

**S5 Sanger confirmed NA12878 chr4_187891282_G_T DNV**
Chromatogram of the DNV, the pink rectangle highlights the DNV position.

**S6 Sanger confirmed NA12878 chr3_77947072_T_A DNV**
Chromatogram of the DNV, the pink rectangle highlights the DNV position.

**S7 Sanger confirmed NA12878 chr8_64304207_C_A DNV**
Chromatogram of the DNV, the pink rectangle highlights the DNV position.

**S8 Sanger confirmed NA12878 chr7_15905632_C_T DNV**
Chromatogram of the DNV, the pink rectangle highlights the DNV position.

**S9 Sanger confirmed NA12878 chr3_151755926_G_T DNV**
Chromatogram of the DNV, the pink rectangle highlights the DNV position.

**S10 Sanger confirmed NA12878 chr6_93124594_T_C DNV**
Chromatogram of the DNV, the pink rectangle highlights the DNV position.

**S11 Sanger confirmed NA12878 chr4_65406505_A_G DNV**
Chromatogram of the DNV, the pink rectangle highlights the DNV position.

**S12 Sanger confirmed NA12878 chr1_247280457_C_A DNV**
Chromatogram of the DNV, the pink rectangle highlights the DNV position.

**S13 Sanger confirmed NA12878 chr8_48494353_G_A DNV**
Chromatogram of the DNV, the pink rectangle highlights the DNV position.

**S14 Sanger confirmed NA12878 chr2_198559835_T_G DNV**
Chromatogram of the DNV, the pink rectangle highlights the DNV position.

**S15 Sanger confirmed NA12878 chr4_5461303_G_A DNV**
Chromatogram of the DNV, the pink rectangle highlights the DNV position.

**S16 Sanger confirmed NA12878 chr12_91353615_T_C DNV**
Chromatogram of the DNV, the pink rectangle highlights the DNV position.

**S17 Sanger confirmed NA12878 chr9_119239892_T_A DNV**
Chromatogram of the DNV, the pink rectangle highlights the DNV position.

**S18 Sanger confirmed NA12878 chr10_109204409_C_T DNV**
Chromatogram of the DNV, the pink rectangle highlights the DNV position.

**S19 Sanger confirmed NA12878 chr6_71729112_T_C DNV**
Chromatogram of the DNV, the pink rectangle highlights the DNV position.

**S20 Sanger confirmed NA12878 chr13_81142986_A_G DNV**
Chromatogram of the DNV, the pink rectangle highlights the DNV position.

**S21 Sanger confirmed NA12878 chr5_106645099_AT_A DNV**
Chromatogram of the DNV, the pink rectangle highlights the DNV position.

**S22 Sanger confirmed NA12878 chr5_123998792_A_C DNV**
Chromatogram of the DNV, the pink rectangle highlights the DNV position.

**S23 Sanger confirmed NA12878 chr19_36641978_C_T DNV**
Chromatogram of the DNV, the pink rectangle highlights the DNV position.

**S24 Sanger confirmed NA12878 chr13_71846473_C_A DNV**
Chromatogram of the DNV, the pink rectangle highlights the DNV position.

**S25 Sanger confirmed NA12878 chr2_155841078_G_A DNV**
Chromatogram of the DNV, the pink rectangle highlights the DNV position.

**S26 Sanger confirmed NA12878 chr20_24046519_G_A DNV**
Chromatogram of the DNV, the pink rectangle highlights the DNV position.

**S27 Sanger confirmed false positive NA12878 chr10_38526918_T_A DNV**
Chromatogram of the DNV, the pink rectangle highlights the DNV position. This variant is a false positive, as there was no alternate allele signal found in NA12878.

**S28 Inconclusive Sanger results for NA12878 chr11_134531608_C_G DNV**
Chromatogram of the DNV, the pink rectangle highlights the DNV position. There is no alternate allele signal on the forward NA12878, but there is a small signal of the alternate allele found in the reverse.

**S29 Partial Sanger confirmation of NA12878 chr21_19977487_C_T DNV**
Chromatogram of the DNV, the pink rectangle highlights the DNV position. We see that NA12878 has an alternate allele signal in both forward and reverse strands. We were unable to obtain the data for the NA12892 reverse strand to fully confirm if the variant is *de novo*.

**S30 Browser shot of ONT sequenced NA12891, NA12892, and NA12878 around DNV chr11_134531608_C_G**
Integrated Genome Browser (IGV) snapshot of the reads around the DNV chr11_134531608_C_G for samples NA12891, NA12892, and NA12878. The boxes show the sequenced reads at the position of the DNV.

**S31 EBV Copy Number Estimation**

A) A distribution of estimated EBV copy number vs total number of DNVs. There was a minor correlation between EBV copy number and total DNVs (p = 2.32e-05, r = 0.17).
B) A violin plot comparing estimated EBV copy number and groups of less than or equal to 100 DNVs and greater than 100 DNVs.

**S32 Phase distributions across autosomal chromosomes**
Distribution of phased DNVs across 6 samples different samples. The blue marks represent paternal parent-of-origin and the red mark represents the maternal parent-of-origin. A) represents a normal distribution, B) represents a clustering of maternal mutations, C), D) and F) represent paternal clustering of mutations, and E) represents a clustering of maternal mutations possible event that could lead to further DNVs.

**S33 CPU vs GPU workflow comparison**

Venn diagrams showing the overlap between detected DNVs for the CPU and GPU version of our workflow on samples A) NA12879 B) monozygotic twin 1 from the SSC cohort and C) twin 2.

**S34 Total counts of variants for DeepVariant and Haplotyepcaller trio calls.**
Distribution of variants by chromosome found in the trios from the 602 1000 Genomes Project families. The total number of variants in blue, and heterozygous variants, in green.

**S35 Total counts of variants for DeepVariant and Haplotyepcaller down-sampled 30x coverage NA12878 trio calls**
Distribution of variants by chromosome found in the trios from the various NA12878 samples, down-sampled to 30x coverage, as well as samples NA12891 and NA12892. The total number of variants in blue, and heterozygous variants, in green.

**S36 SAMtools tview visual validation output example**
An example of how the manual evaluation of the tviews was done. Comparing the first position of each tview, if there was any alternate allele found in the mother or father sample, the tview was denoted as inherited, no matter the position or quality of the variant. If the child had alternate alleles, the DNV would have be validated. The pink rectangles show which column is being examined, with a zoomed in view of the sequences provided.
